# Protoporphyrin-IX nanostructures modulate their protein aggregation ability via differential oxidation and protein binding

**DOI:** 10.1101/2021.01.11.426224

**Authors:** Dhiman Maitra, Benjamin M. Pinsky, Amenah Soherawardy, Haiyan Zheng, Ruma Banerjee, M. Bishr Omary

**Affiliations:** Center for Advanced Biotechnology and Medicine, Rutgers University, Piscataway, NJ 08854; Department of Biological Chemistry, Ann Arbor, MI 48109; University of Michigan Medical School, Ann Arbor, MI 48109

**Author notes:** To whom correspondence should be addressed: Center for Advanced Biotechnology and Medicine, Rutgers University, 679 Hoes Lane W, Piscataway, NJ 08854.

**Keywords:** Porphyrin nanostructure, porphyria, protein aggregation, oxidative stress, photodynamic therapy

## Abstract

Porphyrias are caused by genetic defects in the heme biosynthetic pathway and are associated with accumulation of high levels of porphyrins that become cytotoxic. Porphyrins, due to their amphipathic nature, spontaneously associate into different nanostructures but very little is known about the effect of porphyrin speciation on the cytotoxic effects of porphyrins. Previously we demonstrated the unique ability of fluorescent biological porphyrins, including protoporphyrin IX (PP-IX), to cause organelle selective protein aggregation, which we posit to be a major mechanism by which porphyrins exerts their cytotoxic effect. Herein, we tested the hypothesis that PP-IX-mediated protein aggregation is modulated by different PP-IX nanostructures via a mechanism that depends on their oxidizing potential and protein binding ability. We demonstrate that PP-IX nanostructure formation is reversible in nature, and that nanostructure size modulates consequent protein oxidation and aggregation potential. We also show that albumin, the most abundant serum protein, preferentially binds PP-IX dimers and enhances their oxidizing ability. Additionally, extracellular albumin protects from intracellular porphyrinogenic stress and protein aggregation by acting as a PP-IX sponge. This work highlights the importance of PP-IX speciation in the context of the porphyrias, and offers insights into potential novel therapeutic approaches.

## INTRODUCTION

Protoporphyrin-IX (PP-IX) is nature’s template for several essential biomolecules including heme, chlorophyll, coenzyme F430 (in methanogenic bacteria), and vitamin B12 (1). The PP-IX macrocycle consists of four pyrrole rings connected by methene bridges, and two ionizable propioniate side chains. The highly conjugated macrocycle with 18 π-electrons confers PP-IX with distinct UV-visible absorbance and fluorescence properties (2). Due to its amphiphilic nature, PP-IX tends to aggregate in aqueous solution (3). Indeed, PP-IX has been reported to exist as detergent micellarized monomers (4), dimers (5), tetramers (6), and higher order structures that are >70 kDa in size (4). The spontaneous self-association of PP-IX occurs by a combination of hydrogen-bonding (between the propionate carboxylate), and π-π stacking of the porphine ring (3,6,7). The resulting face-to-face or ‘H-oligomers have the propionate chains of adjacent PP-IX molecules in a ‘head-to-tail’ orientation (3,6,7). These supra-molecular higher order structures offer advantages such as improved responsiveness, reversibility, tunability, biomimicry, modularity, predictability, and adaptiveness, over their constituent monomers (8,9). This self-assembling property of PP-IX and similar compounds are being actively explored for a variety of applications (10–12) and synthetic porphyrin nanostructures for photodynamic therapy (13–15), photovoltaic cells (16–18), photocatalysis (19–23), and for sensor applications (23–25), have been reported. Although porphyrin macromolecular species have been extensively studied, their role in the context of protein aggregation and in the context of porphyrias is not known.

Porphyrias include eight genetic disorders that are caused by errors in any of the eight steps in the heme biosynthetic pathway (26,27). Heme biosynthesis starts in the mitochondrion where aminolevulinic acid (ALA) synthase catalyzes the rate-limiting conversion of glycine and succinyl Co-A to δ-ALA (27–30). Upon entry into the cytosol, ALA is converted in several steps, to the first cyclic tetrapyrrole, uroporphyrinogen, which is subsequently converted to coproporphyrinogen (27–30). Coproporphyrinogen then enters mitochondria, where it is converted to heme via formation of protoporphyrinogen and PP-IX (27–30). Porphyrinogens are colorless, non-fluorescent (31) compounds, which are auto-oxidized to more stable and fluorescent porphyrins (27). Porphyrins are toxic metabolites, and their levels are tightly regulated. In porphyrias, blockages in the heme biosynthetic pathway leads to the accumulation of intermediates, with consequent organ and tissue damage (4,18,25).

Porphyrin-mediated tissue damage is proposed to occur through reactive oxygen species (ROS) generated by type I/II photosensitized reactions of porphyrins (32–34). However, the precise nature of the ROS as well as the specific targets of porphyrin-generated ROS are poorly understood (27). Recent findings demonstrated the unique properties of fluorescent porphyrins to cause organelle-selective protein aggregation though a mechanism that involves a ‘porphyrination-deporphyrination’ cycle (27,35–39). In this cycle, PP-IX binds to target proteins (i.e. porphyrination) and induces localized unfolding and conformational change (40,41). Photosensitization of protein-bound PP-IX leads to the generation of singlet oxygen (^1^O_2_), and oxidation of specific methionines to methionine sulfone or sulfoxide (27,42). Subsequent non-covalent interactions lead to protein-PP-IX lattice-like aggregate formation. During deporphyrination, acidic pH, or high salt extraction of PP-IX leads to disaggregation of the protein aggregates (27,42). We posit that this proteotoxic property of porphyrins is a major mechanism for tissue damage in porphyrias, which involves fluorescent porphyrin accumulation.

In this study, we tested the hypothesis that PP-IX-mediated protein aggregation is modulated by PP-IX speciation into supra-molecular structures, through a mechanism that involves differential oxidizing potential and protein binding.

## RESULTS

### pH-mediated spontaneous and reversible transformations of PP-IX nanostructures

PP-IX speciation results in distinct UV-visible and fluorescence emission signatures (3–5,43). In aqueous solution, pH and ionic strength are the principal modulators of PP-IX speciation (3). We investigated the nature of PP-IX speciation at pH 7.4 (physiological), pH 4.5 (lysosomal) and pH 9, using previously reported spectra for assignments (3–5,43). UV-visible and fluorescence spectra were collected by diluting freshly prepared PP-IX stock solutions in the indicated buffers (Fig.1A,B). In 100 mM HCl, PP-IX exists as monomers, characterized by the sharp Soret band at 409 nm (Fig.1A, Table 1). At pH 4.5, PP-IX monomers form H-aggregates, characterized by a broad Soret band with shoulders at 356 and 466 nm. At pH 9 and in 100 mM NaOH, PP-IX exists exclusively as dimers, as judged from the characteristic Soret peak at 380 nm. Notably, the absorbance spectrum at pH 7.4, shows a combination of features seen at pH 4.5, and 9, displaying a broad Soret band with slightly red shifted shoulders (379 and 469 nm) compared to the pH 4.5 spectrum (Fig1A, Table1). This suggests that at pH 7.4, PP-IX consists of a mixture of higher order aggregates and dimers. In addition to the changes in the Soret band, changes in the Q-bands in 500-700 nm region are seen. In 100 mM HCl, where the four pyrrole nitrogens are expected to be protonated, two Q-bands are observed. The number of Q bands increases to 3 (pH 4.5, 7.4), and 4 (pH 9) as the extent of protonation decreases.

**Table 1:**
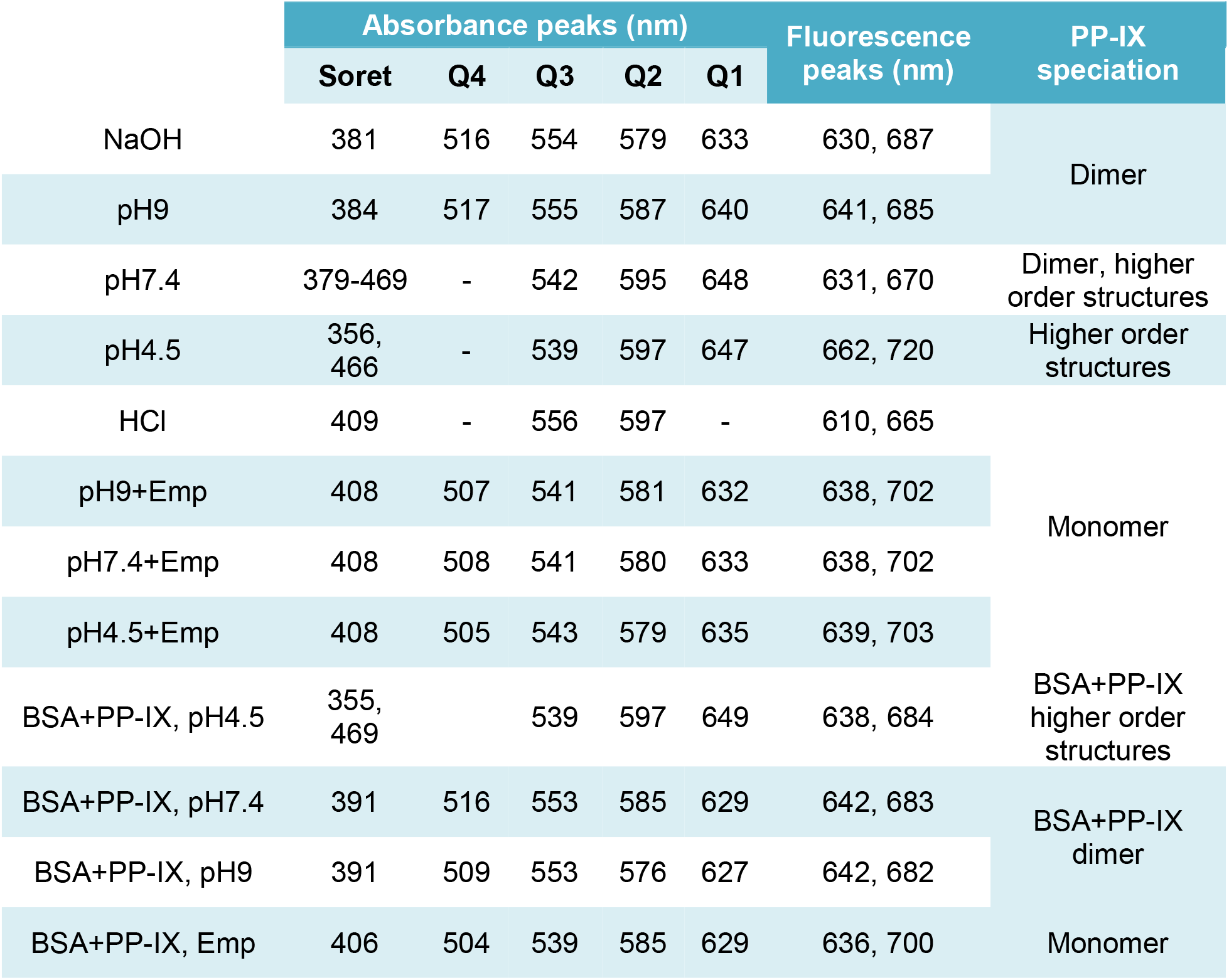
Summary of UV-vis and fluorescence spectral features of different PP-IX nanostructures as shown in Figure 1A, B and E and Figure 4A-D.

**Figure 1:**
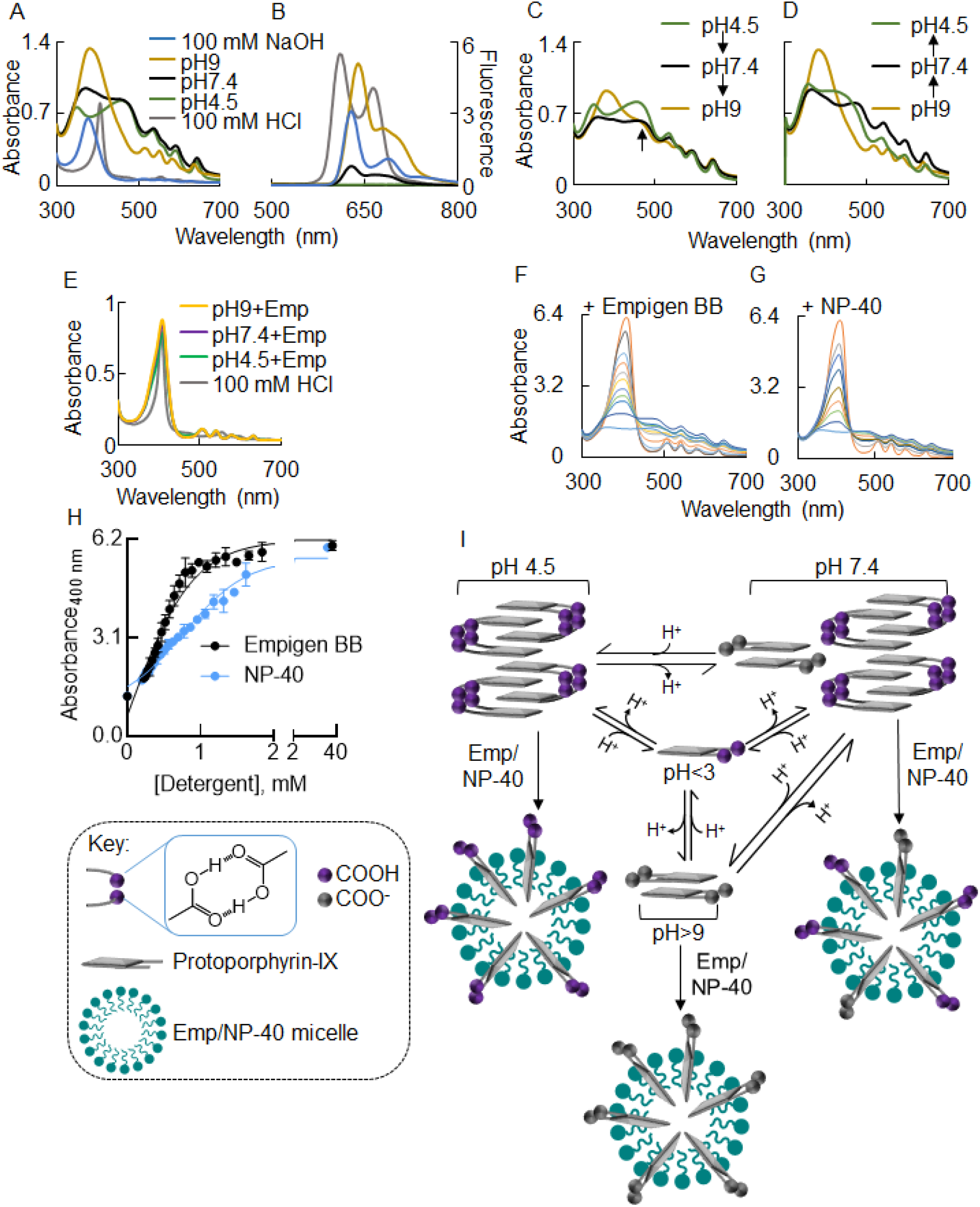
pH-mediated spontaneous and reversible transformations of PP-IX nanostructures. **A)** Absorbance spectra of PP-IX (50 μM) in pH4.5, 7.4, 9, and PP-IX (10 μM) in 100 mM NaOH/HCl. **B)** The PP-IX solutions described in panel A were excited at 375 nm followed by collection of the fluorescence emission spectra from 500 to 800 nm. **C)** Protoporphyrin-IX stock solution was diluted in pH4.5 buffer to a final concentration of 50 μM. The resulting solution was divided into two aliquots, and the pH was then adjusted to 9 or 7.4 using NaOH (5N). Absorbance spectra of the resulting pH9 and pH7.4 solutions were recorded and compared to the absorbance spectra of PP-IX solution at pH4.5. **D)** Protoporphyrin-IX stock solution was diluted using pH9 buffer to a final concentration of 50 μM. The resulting solution was divided into two aliquots, and pH was adjusted to 7.4 and 4.5 using HCl (5N). Absorbance spectra of the resulting pH7.4 and pH4.5 solutions were recorded and compared to the absorbance spectra of PP-IX solution at pH9. **E)** Absorbance spectra of PP-IX (10 μM) in buffers of pH4.5, 7.4 and 9 supplemented with 1% Empigen BB (a zwitterionic detergent; highlighted as ‘Emp’ in panel E), and 100 mM HCl, respectively. **F, G)** Absorbance spectra of PP-IX (10 μM) in 400 mM phosphate buffer (pH7.4), supplemented with increasing concentrations of Empigen BB (panel F) or NP-40 (panel G). The traces shown in panel F corresponds to 0, 0.22, 0.34, 0.38, 0.42, 0.47, 0.52, 0.58, 0.64, 0.98, and 36.76 μM Empigen (bottom to top), respectively. Panel G, shows the absorbance spectra of PP-IX in 400 mM phosphate buffer (pH7.4) with 0, 0.20, 0.33, 0.51, 0.78, 1.06, 1.31, 1.62, 32.41 μM NP-40 (bottom to top), respectively. The absorbance spectra shown in panels A-G are representative of three independent experiments. **H)** Absorbance at 400 nm for the PP-IX solutions described in panels F and G, were plotted as a function of detergent concentration, and the data was fitted to a sigmoidal-dose response curve using GraphPad Prism 8. The data shown is an average of three independent experiments ± standard deviation. **I)** A schematic that summarizes the findings in Figure 1 and highlight the reversible conversion of PP-IX into monomer, dimer and higher order structures as a function of pH and detergent.

The fluorescence emission spectra of the different PP-IX species (Fig.1B) revealed that a significant fluorescence quenching is associated with higher order aggregates of PP-IX. Thus, at pH 4.5, PP-IX displays a minimal fluorescence signal compared to pH 7.4 and 9 (Fig.1B, green trace).

The pH-induced PP-IX speciation is reversible (Fig.1C,D). Thus, the higher order PP-IX structures observed at pH 4.5, convert to a mixture of higher order structures and dimers (pH 7.4), and then dimers (pH 9) as the pH is increased with 5N NaOH (Fig.1C), albeit conversion is incomplete (see shoulder indicated by arrow). The transition in the reverse direction i.e. dimers → higher order structures, was observed when the pH of a PP-IX solution at pH 9 was progressively decreased to 7.4 and 4.5 with 5N HCl (Fig.1D). In earlier studies, we had reported that PP-IX induces protein aggregation in buffers containing detergents (e.g. Empigen BB, NP-40) (36,37,39,42). Given the profound effect of pH on PP-IX speciation, we tested whether detergent plays a role in porphyrin speciation. We found that irrespective of pH, 1% Empigen BB, a non-denaturing, zwitterionic detergent, shifts the equilibrium completely to monomeric PP-IX, mimicking the effect of 100 mM HCl (Fig.1E). To probe the underlying mechanism, a solution of PP-IX (10 μM) in pH7.4 phosphate buffer was supplemented with increasing concentrations of Empigen BB (Fig.1F), or NP-40, a non-ionic non-denaturing detergent, (Fig.1G) and observed the transition from higher order structures/dimers to monomers (Fig.1F-G). The absorbance at 400 nm showed a sigmoidal dependence on detergent concentration, which is characteristic of micelle formation (Fig.1H). Thus detergent-induced PP-IX monomerization appears to be caused by incorporation of monomers into detergent micelles. In summary, we observed that PP-IX reversibly transitions from dimers (at high pH) to a mixture of dimers/higher order structures to higher order structures (at low pH). Detergents and pH values of <3 favor PP-IX monomers (Fig.1I).

### Smaller PP-IX nanostructures have higher oxidizing potential

Since higher order PP-IX structures show profound fluorescence quenching (Fig.1B), we characterized the effect of oligomerization on the quantum yield of PP-IX. As the nanostructure size decreases with increasing pH or with detergent, there is a significant increase in quantum yield (Φ) (Fig. 2A, supplementary Table S2). PP-IX monomers (observed in 1% Emp, pH7.4) exhibit the highest Φ.

**Figure 2:**
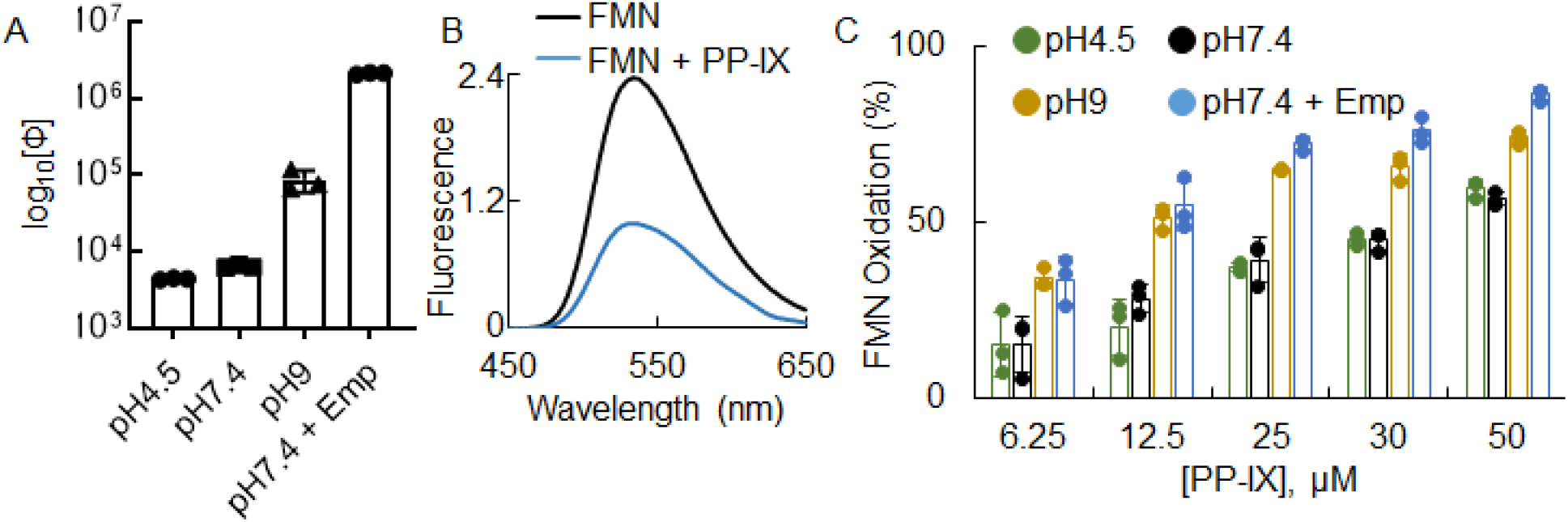
Smaller PP-IX nanostructures have higher oxidizing potential. **A)** Quantum yield of different sized PP-IX nanostructures. The data shown is the average of three independent experiments ± standard deviation. **B)** Fluorescence emission spectra of FMN (21 μM) ± PP-IX (50 μM) at pH7.4 were recorded after exciting the solutions at 400 nm. The data shown is representative of three independent experiments. **C)** PP-IX-mediated FMN oxidation at different pH and PP-IX concentrations. The data shown is the average of three independent experiments ± standard deviation.

Next, we tested whether the size-dependent increase in Φ translates to an increase in PP-IX-mediated photooxidation. For this assay, we examined PP-IX-mediated oxidative destruction using the chromophore FMN (Fig.2B). PP-IX-mediated FMN oxidation increased as a function of PP-IX concentration (Fig.2C). Importantly, the smaller PP-IX structures (monomers and dimers, observed at pH 9 and pH 7.4 + Emp), caused significantly greater FMN oxidation (Fig.2C, supplementary Table S3). In contrast a significant difference in the oxidizing capacity of PP-IX at pH 4.5 and 7.4 was not observed. These data suggest that there are two groups of PP-IX nanostructures with different oxidizing potentials. One group consists of the higher order structures that exist at pH 4.5 and 7.4, while the second group consists of monomers/dimers at pH 9 or in detergent-containing buffer (pH 7.4).

### PP-IX nanostructure size modulates the type and level of PP-IX-mediated protein aggregation

Next, we tested our hypothesis that the increased oxidizing potential of PP-IX dimers/monomers leads to increased protein aggregation. PP-IX-mediated protein aggregation was clearly observed by SDS-PAGE (Fig.3A, compare lanes ± PP-IX). Specifically, the high molecular weight aggregates that failed to enter the resolving gel (Fig. 3A dotted boxes) were more prominent in the PP-IX treated samples as the pH was increased from 4.5 to 9.

**Figure 3:**
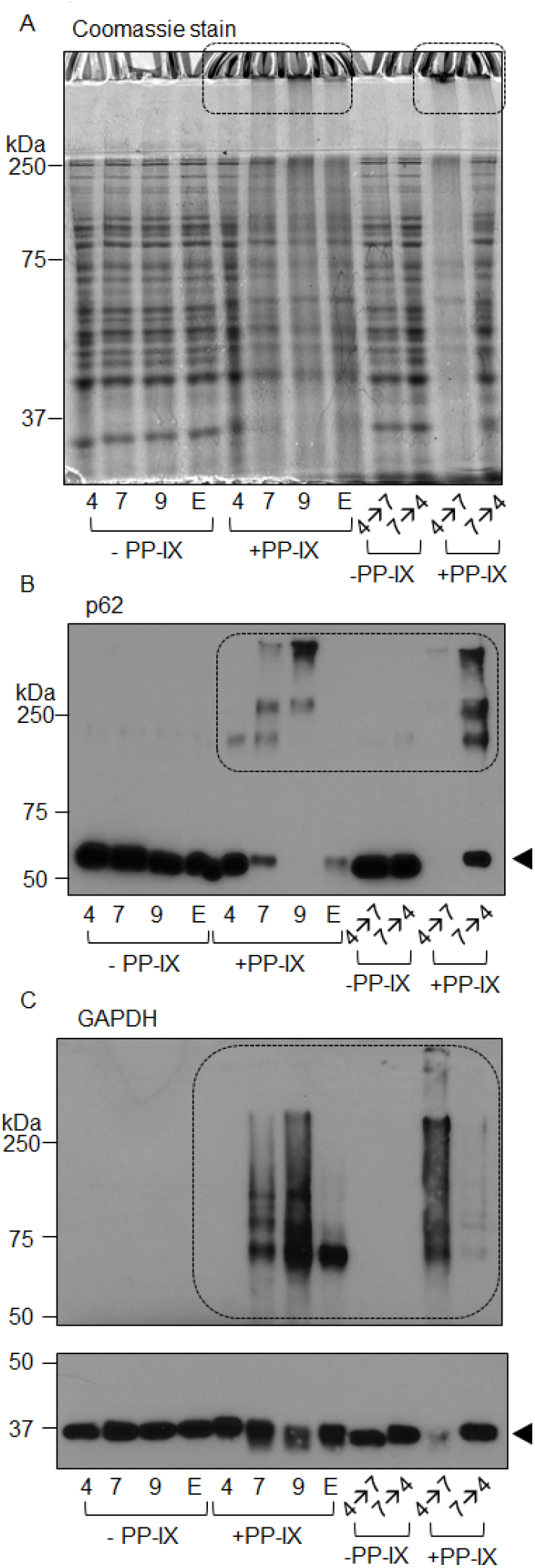
PP-IX nanostructure size modulates the type and quantity of PP-IX-mediated protein aggregation. **A)** The soluble fraction of Huh-7 cell lysate (detailed in Experimental Procedures) was adjusted to 1 mg/mL of protein and treated with 50 μM PP-IX at different pH (‘4’ represents pH4.5, ‘7’ represent pH7.4, ‘9’ corresponds to pH9, and ‘E’ represents pH7.4 + 1% Empigen. In addition, an aliquot of the pH4.5 reaction mixture was adjusted to pH7.4 (‘4→7’) and incubated for further 30 minutes with PP-IX. Similarly, the pH7.4 reaction mixture was adjusted to pH4.5 (‘7→4’). After treatment, the reaction mixtures were quenched with the addition of reducing SDS-PAGE sample buffer. For each lane,10 μg of protein was separated by SDS-PAGE and stained with Coomassie blue. **B,C)** Reaction mixtures described in panel A were separated by SDS-PAGE, transferred to PVDF membranes then blotted with antibodies to p62 (B) and GAPDH (C). Monomer bands are marked with an *arrowhead*, and where shown separately (panel C), the monomer blot (lower panel) was exposed for 5 seconds while the HMW aggregate (marked as HMW agg) was exposed for 15 minutes. The data shown is representative of three independent experiments.

Two soluble proteins, sequestome 1 (p62) and glyceraldehyde 3-phosphate dehydrogenase (GAPDH) also exhibited a dramatic increase in aggregation as the pH increases (Fig.3B,C, dotted boxes) and a corresponding loss of monomers at pH 9 (Fig.3B,C, *arrowhead*). PP-IX treatment caused three distinct tiers of high molecular weight aggregates in p62 (Fig.3B, dotted box). Decreasing the size of PP-IX nanostructures (via a change from pH4.5 → pH9), caused formation of heavier (with decreased electrophoretic mobility) high molecular weight aggregate. Using pH7.4+1% Empigen, we did not observe any high molecular weight aggregate for p62, although there was an evident loss of monomer (Fig.3B, lane ‘E’+PP-IX, *arrowhead*). We attribute the extensive loss of monomer of p62 to the loss of antibody reactivity to protein oxidation and/or aggregation following PP-IX treatment as described previously for lamin B1 (39), cyclin B1 and cdk4, (42). For GAPDH we did not observe a pronounced loss of monomer, (Fig.3D, *arrowhead*). In terms of high molecular weight oligomers, GAPDH showed the trend: pH9> pH7.4> 1% Emp>> pH4.5 (Fig.3D, dotted box). Notably, in contrast to p62, smaller sized PP-IX nanostructures [e.g. dimers (pH9), and monomers (1% Empigen)] lead to formation of smaller-sized high molecular weight structures (Fig3D, dotted box). In summary these results highlight the qualitative and quantitative heterogeneity in protein aggregation depending on the speciation of the PP-IX nanostructures.

### Reversing PP-IX speciation reverses protein aggregation

Since the PP-IX nanostructures are interconvertible (Fig.1C,D), we hypothesized that PP-IX-mediated protein aggregation can be reversed by changing the associated PP-IX nanostructure. To test the hypothesis, we titrated PP-IX in pH 4.5 and 7.4 reaction mixtures to pH 7.4 and 4.5, respectively (Fig.3B,C, lanes 4→7 and 7→4). Notably, reversing the pH from pH 7.4 to 4.5, increased the population of unaggregated p62 and GAPDH (Fig. 3B,C), which we interpret as disaggregation due to loss of porphyrin binding. The opposite effect was observed upon changing the pH of the reaction mixture from 4.5 to 7.4.

### BSA disrupts PP-IX nanostructures by preferentially binding to PP-IX dimers

We tested how PP-IX speciation modulates its binding to fatty acid-free BSA, the most abundant serum protein which has been reported to bind PP-IX (44,45). Incubation of BSA (250 μM) with PP-IX (25 μM) at pH 4.5, 7.4 and 9, increased the absorbance and fluorescence emission intensity of PP-IX, (Fig. 4A-C). In contrast, the BSA-PP-IX mixture in 1% Empigen showed no spectral changes in (Fig.4D), indicating with the absence of binding. At pH 4.5, PP-IX binding to BSA increased both the absorbance and fluorescence intensity but did not change the absorbance maxima or peak shape (Fig.4A, Table1). At pH 7.4, a dramatic shift in the Soret peak was observed in addition to increased intensity (Fig. 4B, Table1), indicating preferential binding of BSA to PP-IX dimers. The increased resolution of the Q-bands from three to four in the presence of BSA (Fig.4B, main panel and inset, Table1), suggested increased pyrrole ring nitrogen deprotonation and/or decreased mobility due to PP-IX binding to BSA. Additionally, at pH7.4 and 9 we also observed a blue shift of Q1/Q2 bands (Fig. 4C, Table1). At pH 9, an increase in intensity was accompanied by a slight red shift in the Soret peak (Fig. 4C, Table1). But Notably, the absorbance spectra of the BSA-PP-IX complex at pH 7.4 versus 9 were indistinguishable (Table 1 and Fig.S1), indicating that the same dimeric PP-IX was bound. Albumin shows perturbations of secondary structure at acidic pH, e.g. 2.4% decrease in percent helix content at pH4 versus pH7 (46). Our data suggests that despite the pH-induced structural perturbation, BSA binds PP-IX higher order nanostructures at pH4.5. In summary we conclude that: (i) there are two classes of PP-IX binding sites on BSA, one for PP-IX higher order nanostructures and a second for PP-IX dimers; ii) BSA binds preferentially to PP-IX dimers at physiological pH; and iii) BSA does not bind to PP-IX monomers.

**Figure 4:**
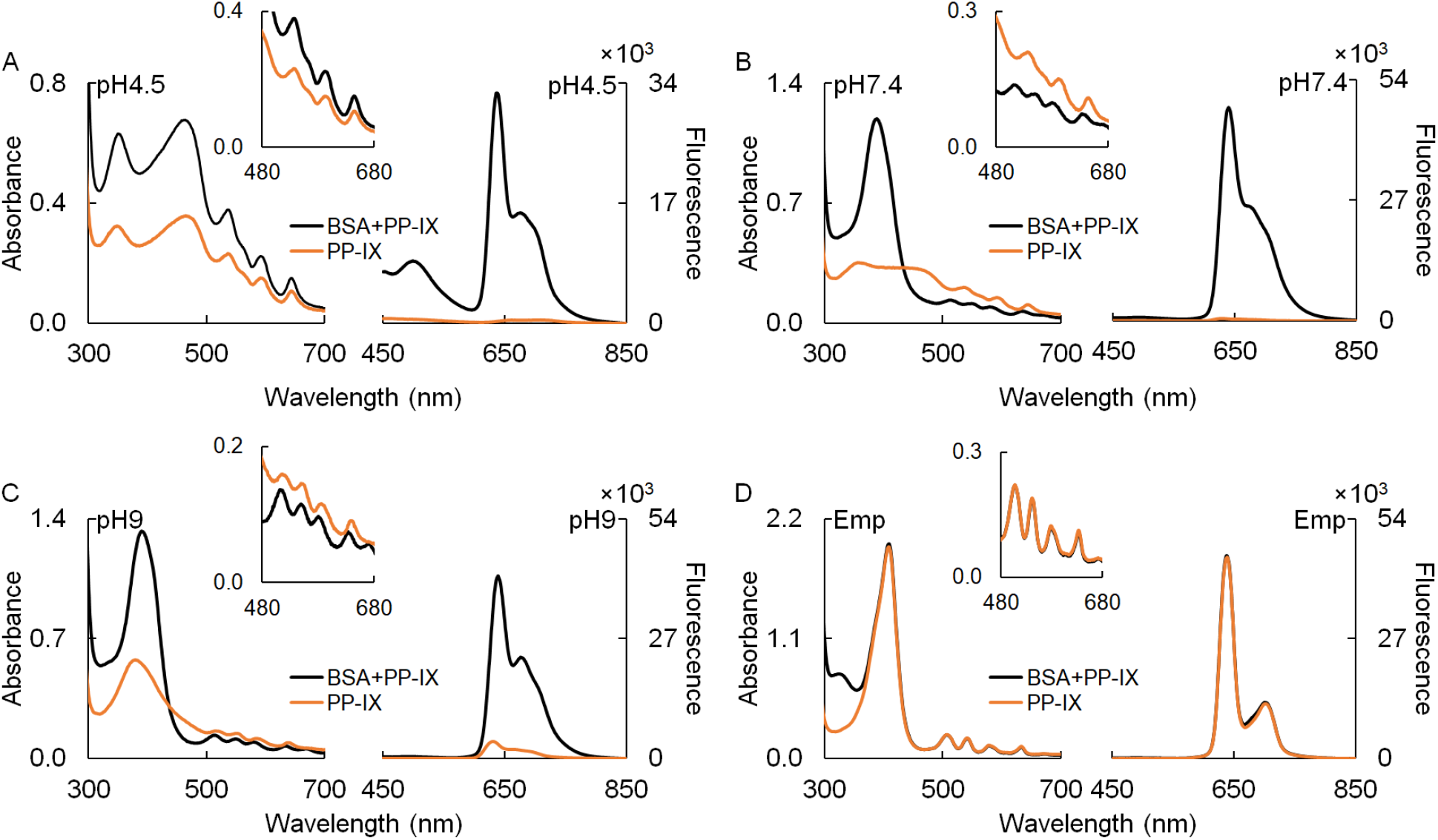
BSA breaks down higher order PP-IX nanostructures by preferentially binding to PP-IX dimers. PP-IX (25 μM) was incubated with BSA (250 μM) at pH 4.5 (panel A), pH7.4 (panel B), pH9 (panel C), and with 1% Empigen (Emp) at pH7.4 (panel D). After a 30-minute incubation, the absorbance and fluorescence (excitation 400 nm) spectra were collected. The insets show a zoomed portion of the absorbance spectra between 480-680 nm to highlight the changes in the Q-band region. The data shown is representative of three independent experiments.

### Differential binding affinity of BSA to different PP-IX nanostructures

We used intrinsic fluorescence of BSA to monitor conformational changes induced by PP-IX binding and estimate the dissociation constant for the different PP-IX nanostructures. Since there is some pH-induced difference in the secondary structure of albumin (46), we examined the intrinsic fluorescence of BSA at pH 4.5, 7.4 and 9. The intensity of the fluorescence emission was sensitive to pH and a small red-shift was observed at pH 7.4 (Fig.5A, inset). When BSA (0.5 μM) was incubated with increasing concentration of PP-IX (0-50 μM), two types of changes were observed: a) a concentration-dependent decrease in the emission intensity (Fig. 5B-E), b) concentration dependent blue shift (Fig. 5B-F). We attribute the blue shift to the increased hydrophobicity in the environment upon PP-IX binding, with subsequent oxidative modification of aromatic residues leading to loss of fluorescence. At pH7.4 (where a mixture of dimers and higher order oligomers exist), the change in fluorescence intensity at 347 nm and the change in emission maximum showed a biphasic dependence on PP-IX concentration (Fig.5E,F). The slope of the steeper part of the curve at pH 7.4 matches that at pH 9, while the second phase matches tat at pH 4.5. This suggests that at low PP-IX concentration, dimers bind to the high affinity dimer binding site on BSA; while at higher IX concentrations, the low affinity ‘higher order structure’ binding site is occupied. To further assess this model, we analyzed the fluorescence quenching data. Only at pH 7.4 two phases were seen, consistent with two classes of binding sites (Fig. 6A-C). The number of binding sites was calculated from the slopes (Table 2), and the dissociation constant (*K*_D_) was determined by plotting F_max_-F versus PP-IX concentration (Fig. 6D). This analysis revealed two high affinity dimer binding sites and two low-affinity higher order nanostructure binding sites per BSA molecule at pH 7.4 (Table 2). Depending on the pH, the cooperativity of PP-IX binding to BSA changed.

**Figure 5:**
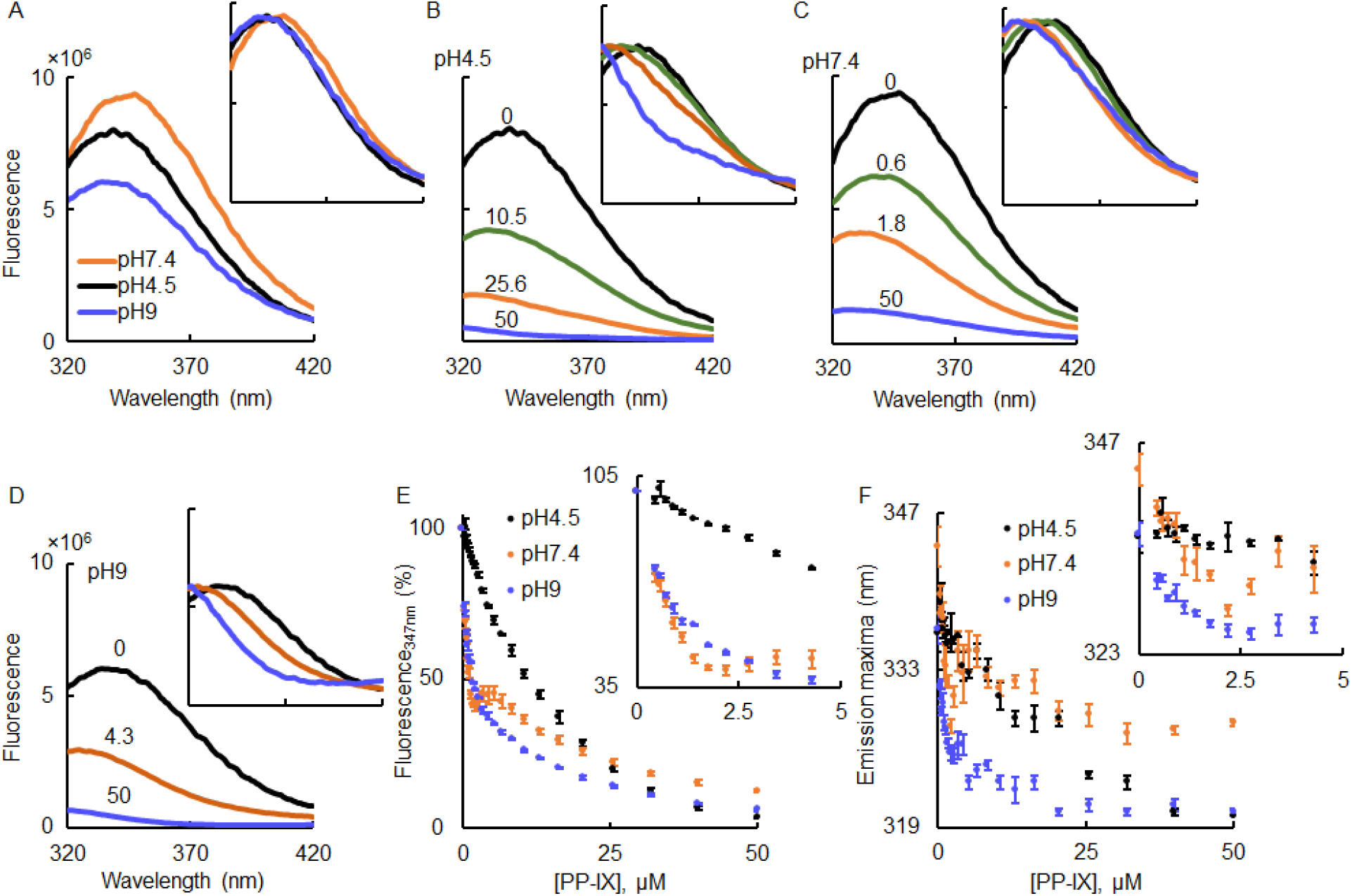
Protoporphyrin-IX binding to BSA leads to a blue shift and loss of BSA intrinsic fluorescence. **A)** BSA (0.5 μM) at different pH, was excited at 280 nm followed by collection of the fluorescence emission from 320 to 420 nm. Inset shows the same spectra normalized to pH7.4 emission maxima. **B-D)** BSA (0.5 μM) was incubated with PP-IX (0-50 μM) for 30 minutes, and emission spectra were recorded after exciting the sample at 280 nm. The numbers above the spectral traces represent PP-IX concentration in μM. Inset shows the same spectra normalized to BSA alone sample emission maxima. The data shown in panels A-D are representative of three independent experiments. **E-F)** Change in BSA fluorescence emission at 347 nm (panel E), and change in emission maxima of BSA (panel F) as a function of PP-IX concentration at the indicated pH. For panel E, the fluorescence emission for ‘BSA alone’ was normalized to 100%. Inset shows the zoomed initial part (0-5 μM PP-IX) of the curve. The data shown is an average of three independent experiments ± standard deviation.

**Figure 6:**
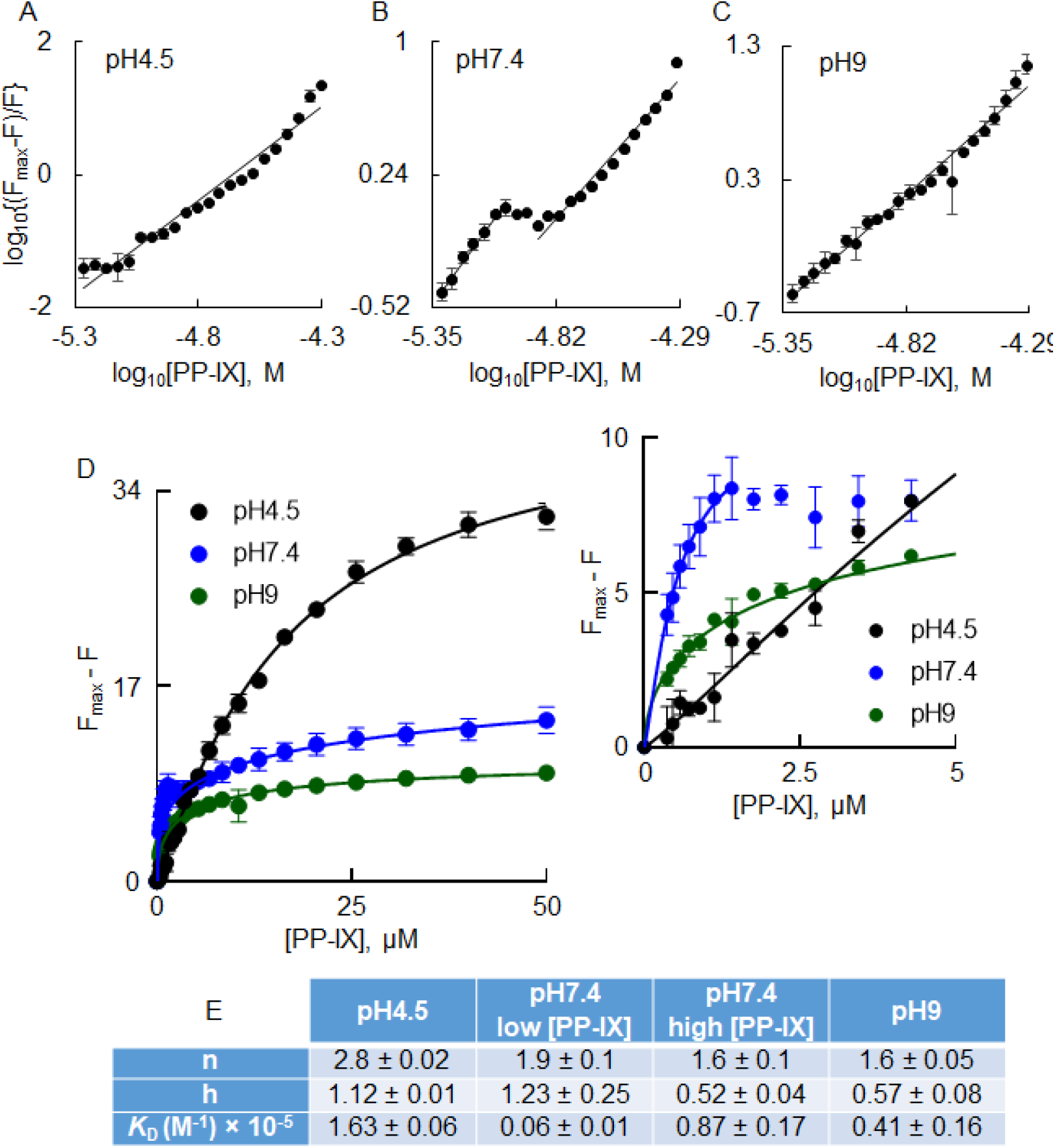
Fluorescence quenching analyses reveal multiple PP-IX binding sites on BSA. Increasing PP-IX concentrations (0-50 μM) were incubated with BSA (0.5 μM) for 30 minutes (pH 4.5, 7.4 or 9). After incubation, quenching of the intrinsic fluorescence of BSA was analyzed by measuring BSA’s florescence emission at 347 nm after exciting at 280 nm. **A-C)** Double log plot of log_10_[(F_max_-F)/F] versus log_10_[PP-IX], at the indicated pH. **D)** Plot of F_max_-F as a function of PP-IX concentration. BSA (0.5 μM) was incubated with 0-50 μM of PP-IX for 30 minutes at the indicated pH. The smooth lines show the fitting of the data to a nonlinear regression model of saturation binding – ‘specific binding with Hill Slope’ equation in GraphPad Prism 8. The inset of panel D shows a zoomed-in portion of the initial part of the curve in order to highlight the binding profile from 0-5 μM PP-IX. The data shown is the average of three independent experiments ± standard deviation. E) Binding parameters for BSA-PP-IX interaction. n, number of binding sites; h, Hill slope; *K*_D_, dissociation constant. Data shown are average of three independent experiments ± standard deviation.

### The BSA and PP-IX complex does not dissociate after urea-induced denaturation

To test the specificity of BSA/PP-IX interaction, we examined the effect of BSA denaturation on PP-IX binding. We denatured BSA with guanidine-HCl or urea, before or after PP-IX binding. Adding PP-IX to either denatured BSA prevented PP-IX binding, as judged by the lack of absorbance or fluorescence changes (Fig. 7A,B). On the other hand, when BSA was denatured after PP-IX treatment we observed a different trend. In the presence of guanidine hydrochloride (Fig.7 C), but not in the presence of urea (Fig.7D), PP-IX was released from BSA. Guanidine-HCl purportedly affects α-helices, while urea affects β-sheets (47). Since albumin is predominantly (64%) α-helical (48), we conclude that guanidine-HCl in more efficient at disrupting its secondary structure.

**Figure 7:**
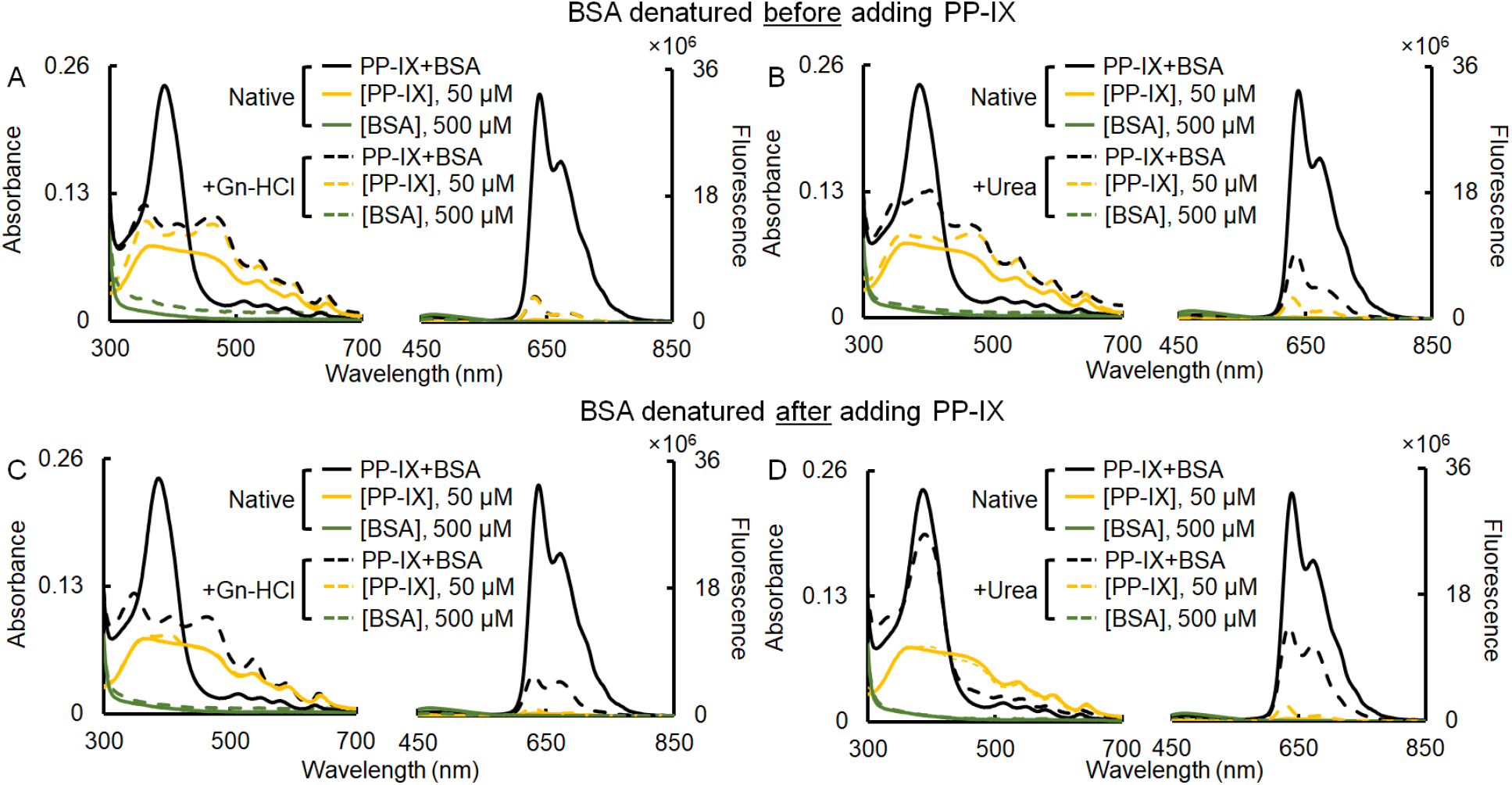
The BSA+PP-IX complex does not dissociate after urea induced denaturation. BSA (500 μM) was incubated with PP-IX (50 μM) (solid lines, ‘Native’). Additionally, BSA and PP-IX solutions were prepared in guanidine (‘+Gn-HCl’, panel A), and urea (‘+Urea’, panel B), respectively. In a parallel set of experiments, ‘Native’ BSA, PP-IX and BSA+PP-IX mixtures were denatured, with guanidine (panel C) or urea (panel D), respectively. The final concentration of Gn-HCl was 6.8 M, and urea was 7.2 M in all cases. Absorbance and fluorescence (excitation at 400 nm) spectra were collected for all the samples. The data shown is representative of three independent experiments.

### PP-IX binding to BSA, enhances its oxidizing ability

When increasing concentrations of BSA (0-500 μM) were incubated with a fixed concentration of PP-IX (25 μM), we observed a concentration-dependent increase in the PP-IX Φ (Fig.8A). To test whether the oxidizing capacity of PP-IX is enhanced upon binding to BSA, the FMN oxidation assay was used. Incubating PP-IX (5 μM) with 0-5 μM BSA caused a concentration-dependent increase in PP-IX absorbance (Fig.8B). FMN oxidation was enhanced in the PP-IX+BSA mixture compared to PPI-IX alone (Fig.8C), and this was observed at all BSA concentrations (Fig. 8D). At higher BSA concentrations, there was a slight decrease in FMN oxidation, suggesting the ability of BSA to act as a sink for PP-IX generated oxidants.

**Figure 8:**
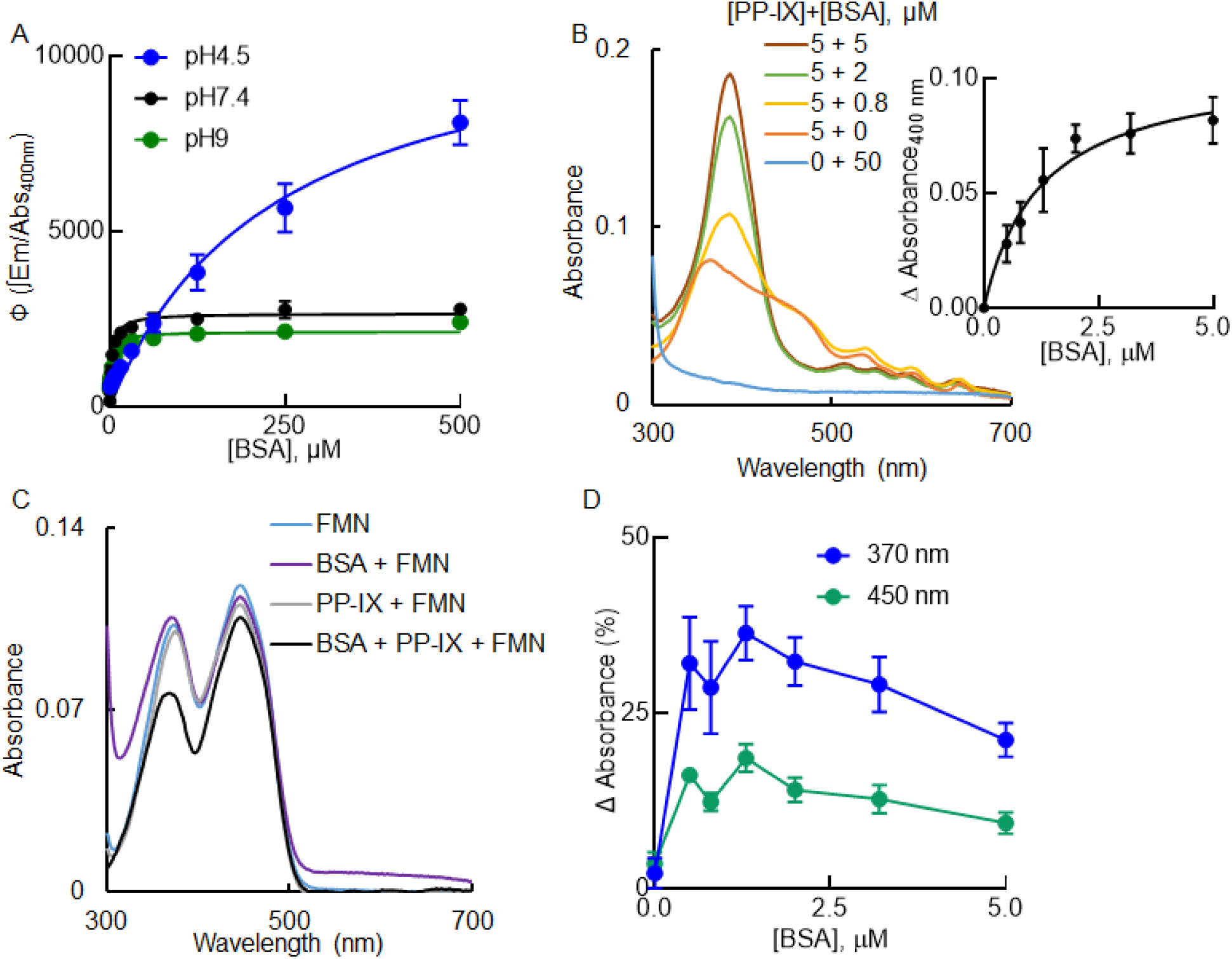
Protoporphyrin-IX binding to BSA enhances the oxidizing ability of PP-IX. **A)** PP-IX (25 μM) was incubated with increasing concentrations of BSA, and the quantum yield (Φ) of PP-IX was plotted as a function of BSA concentration. The data was fitted to a hyperbola (smooth line) using GraphPad Prism 8. **B)** Absorbance spectra of different PP-IX + BSA reaction mixtures. Inset shows the difference in absorbance of PP-IX (calculated from the relative increase in PP-IX absorbance after adding BSA) plotted as a function of BSA concentration. The data was fitted to a hyperbola (smooth line) using GraphPad Prism 8. **C)** Absorbance spectra of FMN (22 μM) alone, BSA (50 μM) + FMN (22 μM), PP-IX (5 μM) + FMN, and BSA (5 μM) + PP-IX (5 μM) + FMN (22 μM). **D)** Percent change in the absorbance at 370 and 450 nm of the different BSA + FMN + PP-IX mixtures with respect to FMN was plotted as a function of BSA concentration. Panels B and C are representative of three independent experiments, and panels A, B (inset) and D show the average of three independent experiments ± standard deviation.

### PP-IX binding does not cause BSA aggregation

Based on the ability of PP-IX to induce protein aggregation [(Fig.3 and (35,36,39,42)], we tested its effect on BSA. PP-IX migrates differently in a pH dependent fashion. Higher order PP-IX nanostructures (at pH 4.5) are more prominent at the top of the gel, particularly under non-reducing conditions (Fig. 9A,D). The band corresponding to the BSA monomer became more diffuse, in the presence of PP-IX, particularly as the pH increased (Fig. 9B,E, *arrowhead*). This is consistent with increased BSA oxidation due to predominance of PP-IX dimers. Importantly, despite PP-IX induced BSA oxidation, there was no evidence of significant aggregation even when the gel was overloaded (Fig. 9C,F).

**Figure 9:**
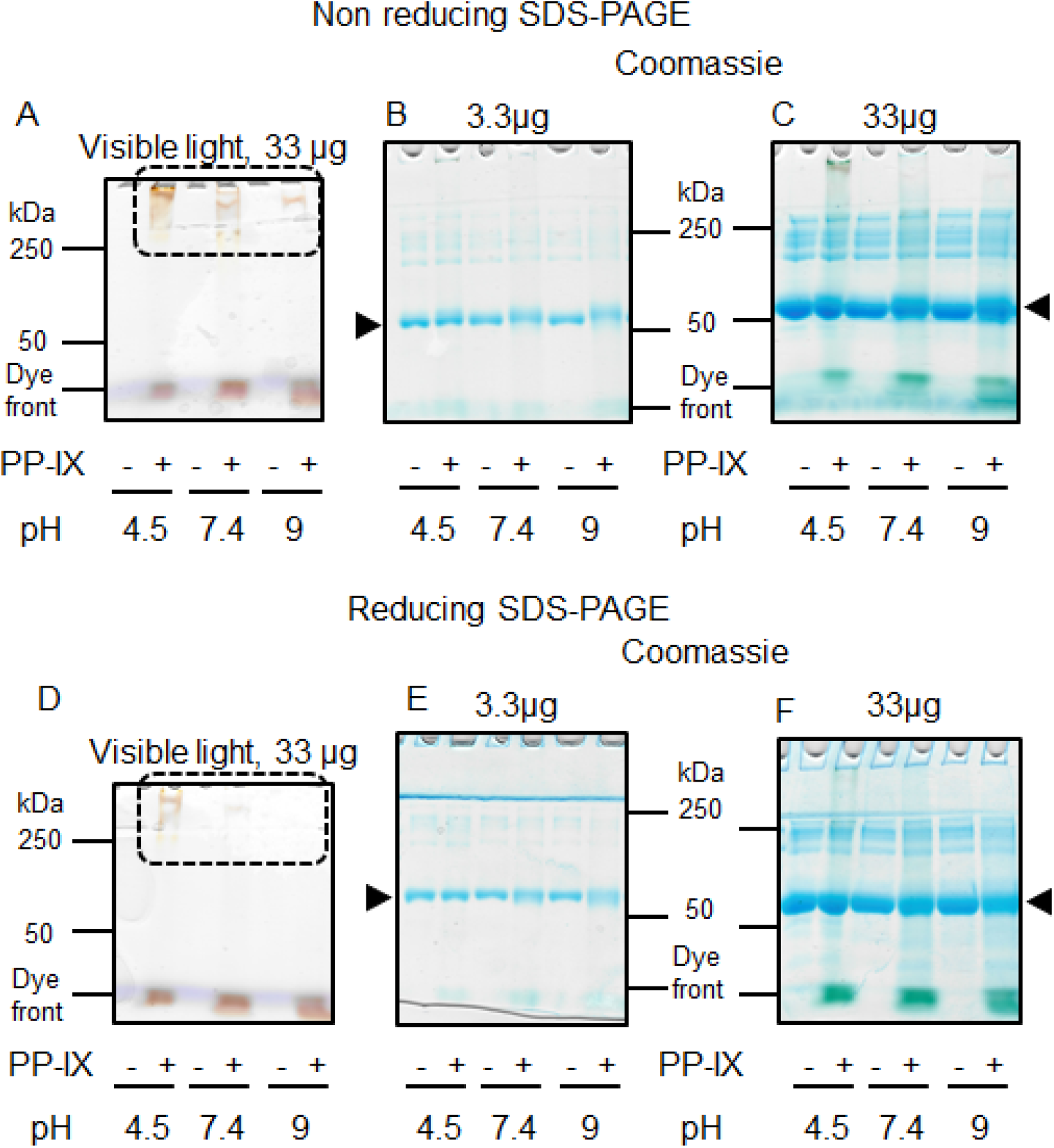
PP-IX binding does not cause BSA to aggregate. BSA (10 μM) was treated with PP-IX (500 μM) for 30 minutes at the indicated pH. The incubation was stopped by adding non-reducing (panels A-C) or reducing (panels D-F) SDS-PAGE sample buffer. After SDS-PAGE, the gels were scanned under visible light (to visualize PP-IX, panels A and D), then stained with Coomassie (panels B, C, E, F) to visualize proteins. The BSA monomer band is marked with an *arrowhead*. The data shown is representative of three independent experiments.

### Unique BSA oxidation signatures based on different PP-IX nanostructures

We utilized LC-MS/MS to identify the site and type of PP-IX-mediated BSA oxidation. Since BSA shows some background oxidation, we filtered our data set and focused on oxidized residues with increased relative abundance following PP-IX treatment in three independent experimental repeats. Over 80% of oxidized residues identified were present only in PP-IX treated samples (Fig.10, boxes with black circles). As expected from the loss of BSA fluorescence emission (Fig. 5), PP-IX-led to oxidation at several tyrosine. In addition, we observed abundant protein carbonyls (+14), but no methionine oxidation (Fig.10). By contrast, methionine oxidation are abundant in other aggregated proteins including keratins and lamins (42).

**Figure 10:**
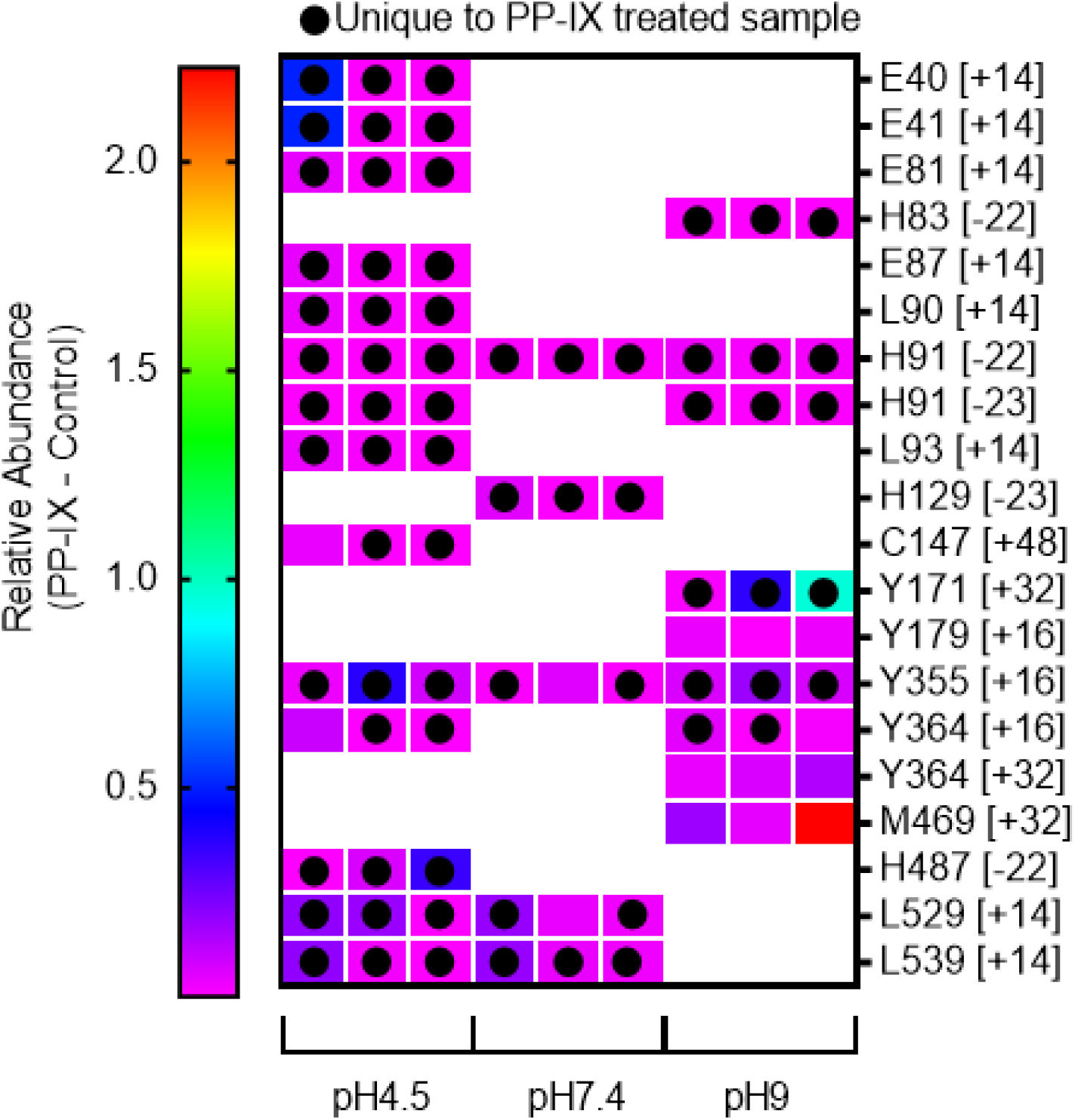
Unique BSA oxidation signatures caused by different PP-IX nanostructures. BSA (10 μM) was treated with PP-IX (500 μM) for 30 minutes at the indicated pH. The reaction mix was stopped by extracting PP-IX followed by analysis of the extract by LC-MS/MS, as detailed in *Experimental procedures*. The heat map shows the relative abundance of the peptides that span the oxidized residues. The amino acid residue (one letter abbreviations) and the type of oxidation (denoted as change in mass; see Supplementary Table S1 for details regarding the type of oxidation) is shown on the right side of the panel. Empty cells within the panel denote residues for which no oxidation was detected using our experimental conditions. Results from three independent experiments are shown.

While some residues (e.g., His-91, Tyr-355) were oxidized at all three pHs, the oxidation signatures showed some pH sensitive differences (Fig.10). All but one (Met-469) of the identified oxidation sites is conserved among bovine, mouse and human albumin (Met-469 is substituted by lysine in murine albumin). We propose that an overlap in the binding sites at different pHs is reflected by the ubiquitously oxidized residues, while residues that are oxidized only at pH 4.5 or 9 report on different sites and/or their level of occupancy (Fig.11).

**Figure 11:**
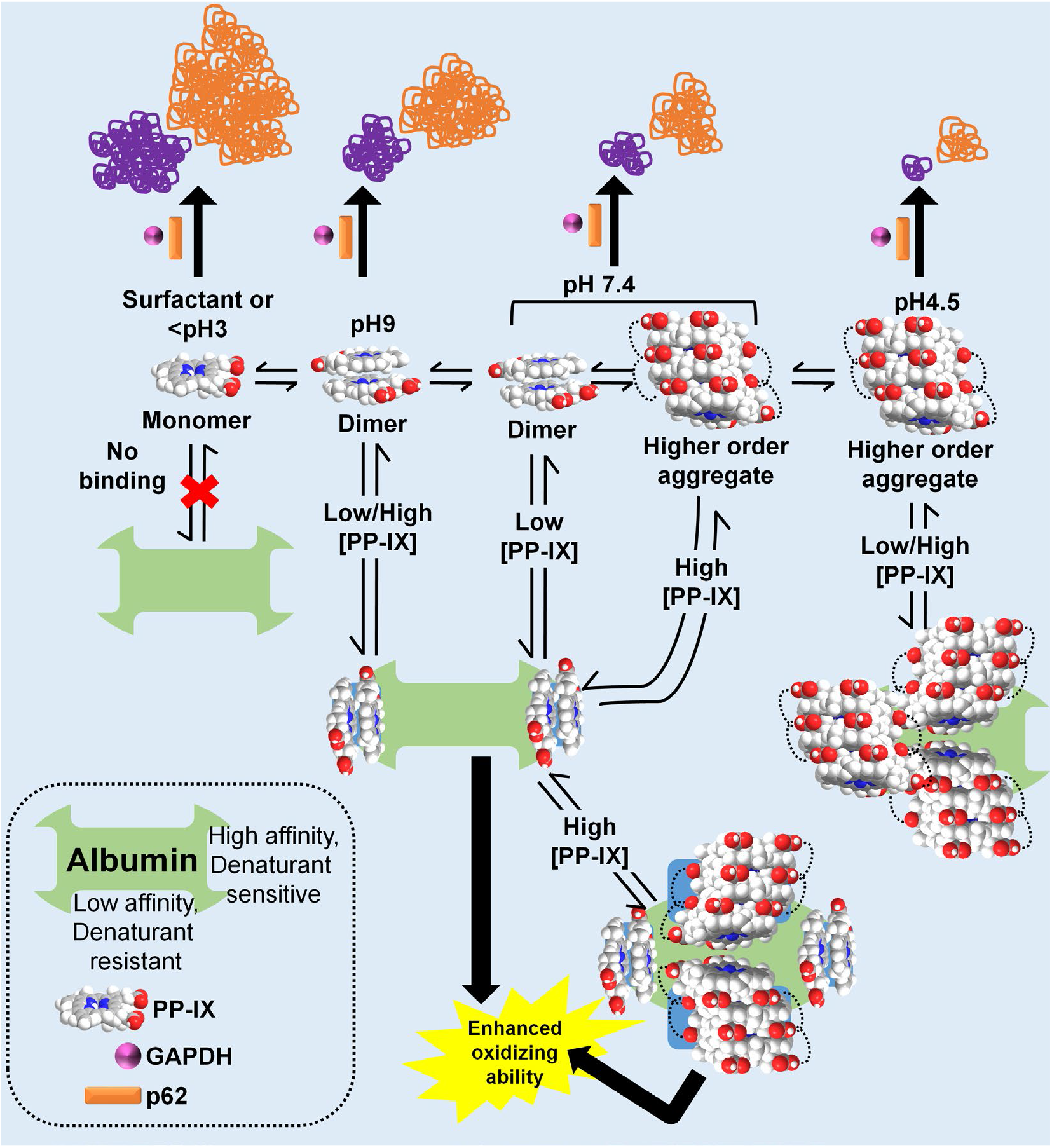
The model summarizes the effect of PP-IX nanostructures in PP-IX-mediated protein aggregation, and the modulation of PP-IX speciation by albumin.

### BSA protects against intracellular porphyrin accumulation and protein aggregation after ALA+DFO treatment

Given the ability of BSA to bind PP-IX, we hypothesized that BSA modulates porphyrin-mediated protein aggregation by acting as an extracellular PP-IX sponge. To test this hypothesis, we treated HuH-7 cells with δ-aminolevulinic acid (ALA)+deferoxamine (DFO) with or without BSA added to the culture medium. ALA+DFO resulted in intracellular accumulation of PP-IX and Copro (Fig.12A,C), as well as their extracellular release (Fig.12B,D). When the medium was supplemented with 5 mg/ml BSA, significantly less porphyrin accumulated inside cells, while more porphyrin was released (Fig. 12C). We attribute this to the ability of BSA to bind PP-IX and shift the equilibrium of PP-IX secretion.

**Figure 12:**
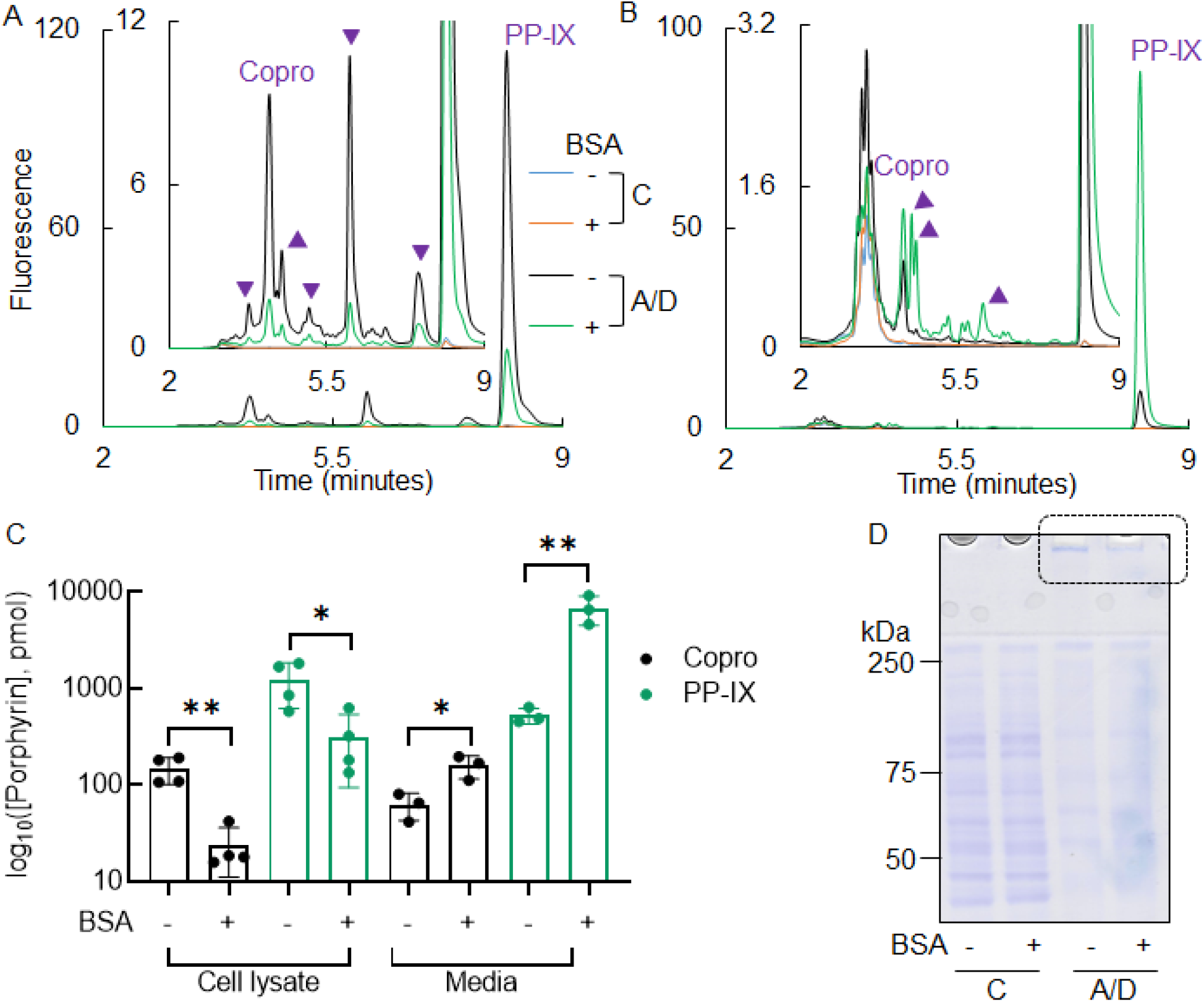
BSA decreases intracellular porphyrin accumulation after ALA+DFO treatment of HuH-7 cells. HuH-7 cells were treated with ALA+DFO for 16h in serum-free media (supplemented with 5 mg/mL BSA where indicated). Porphyrins were then analyzed in the cells and the media by UPLC. **A-B)** UPLC chromatograms show the amounts of fluorescent porphyrins inside the cells (A), and in the media (B). Copro and PP-IX peaks are labelled, and arrowheads indicate unknown fluorescent products that are formed after ALA+DFO treatment. **C)** Quantification of Copro and PP-IX levels calculated from the area under the curve of the fluorescent peaks shown in panel A and B. The data shown are average of three independent experiments with error bars representing the standard error of measurement. Statistical significance was determined using an unpaired t test (2-tailed). *P < 0.05, **P < 0.01 and denotes comparison with control. **D)** Proteins from the collected cells were separated by reducing SDS-PAGE and visualized by Coomassie staining (dotted box highlights the stained aggregates that did not enter the gel). The data shown in panels A-B and D are representative of three independent experiments.

ALA+DFO treated cells with BSA in the medium also had a lower concentration of high molecular weigth protein aggregates as seen by Coomassie blue staining (Fig. 12D, dotted box). We also tested of several specific proteins including cytosolic (K8 and K18), endoplasmic reticulum (BiP and PDI), nuclear (lamins A/C and B1), and proteins involved in the degradation and clearance pathway (p62 and ubiquitin). BiP is an ER-specific HSP70 isoform and a marker for the unfolded protein response (49), and is upregulated after PP-IX induced stress (39,42). BiP is also induced when cells are treated with ALA+DFO in absence of BSA (Fig.13). Notably, all tested proteins were protected by BSA against ALA+DFO induced loss of the monomeric form and aggregation (Fig.13). We posit that albumin by acting as a ‘PP-IX sponge’, protects against intracellular porphyrin accumulation and subsequent aggregation (Fig.14).

**Figure 13:**
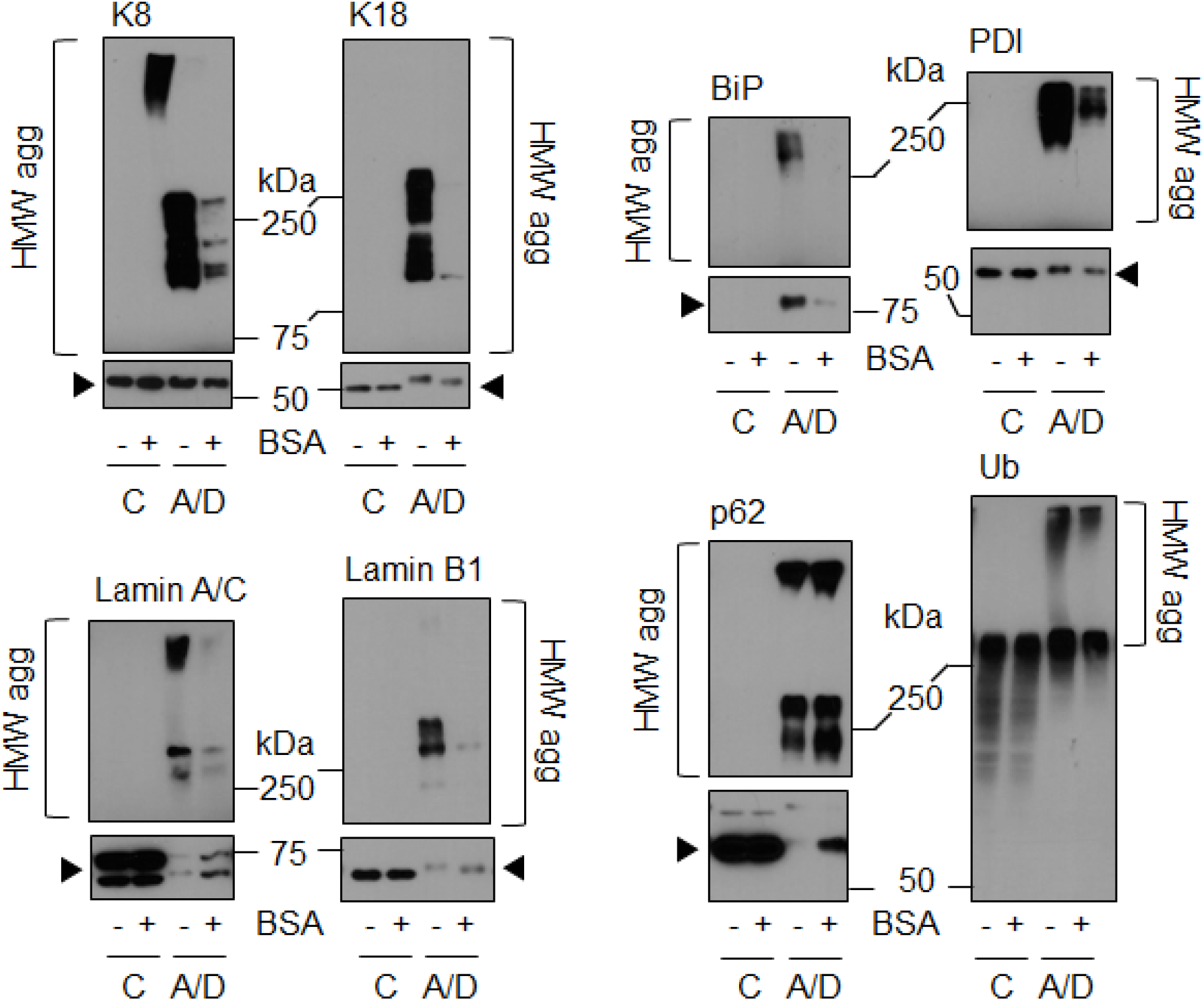
BSA protects from ALA+DFO mediated cellular protein aggregation. Protein from the HuH-7 cells experiment described in Fig.12 were separated in reducing SDS-PAGE, then immunoblotted with antibodies to the indicated antigens. Monomer bands are marked with an *arrowhead* and, where shown separately, the monomer blots (lower panels) were exposed for 5 seconds, while the HMW aggregate blots were exposed for 15 minutes. The data shown is representative of three independent experiments.

**Figure 14:**
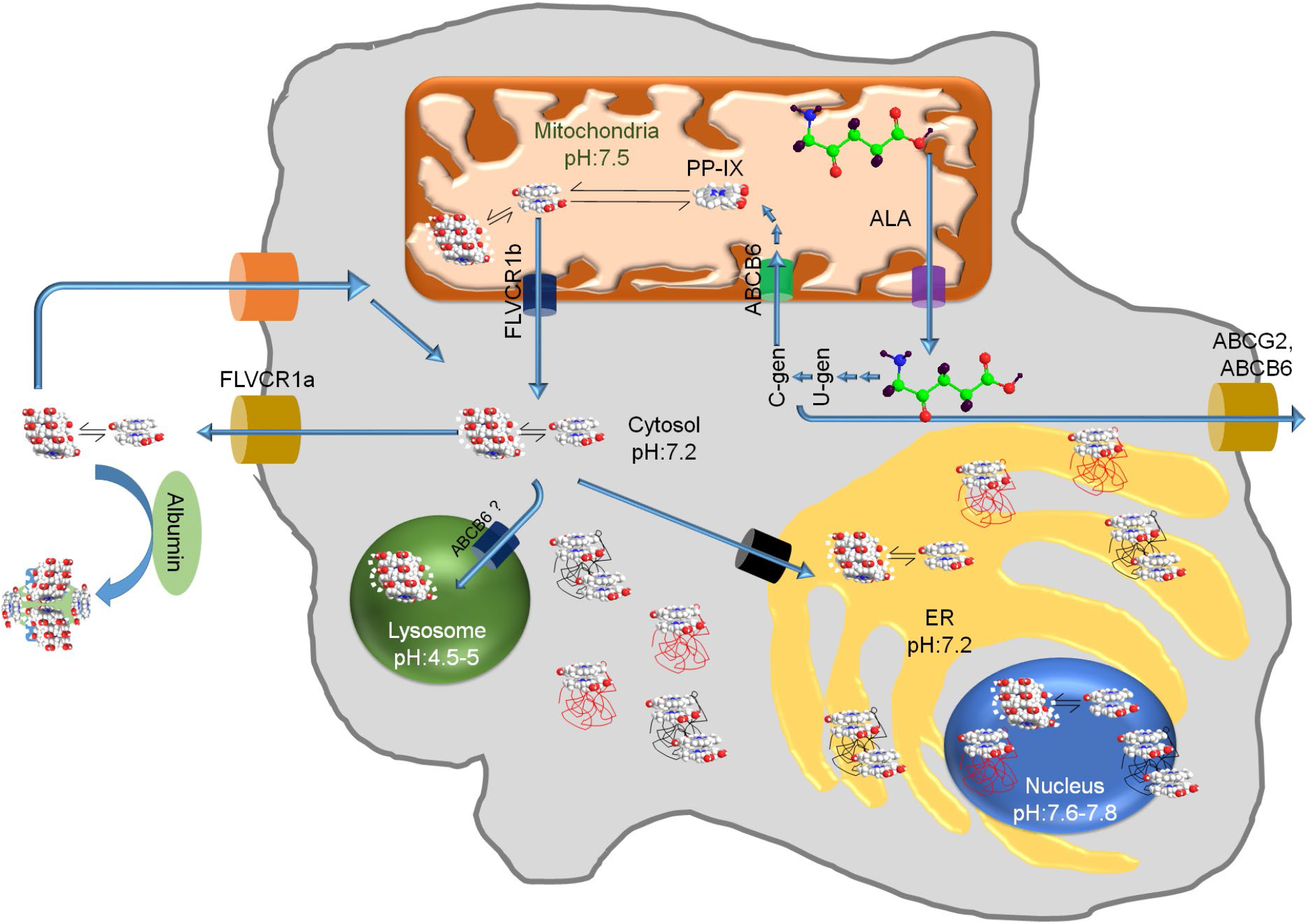
A model for how albumin acts as an extracellular trap for porphyrins, thereby modulating porphyrin-mediated proteotoxicity and cell damage.

## DISCUSSION

### Targeting PP-IX speciation as a potential therapeutic strategy

Porphyrin-mediated protein aggregation plays a major role in porphyric tissue damage (27,39,42). PP-IX oligomerization causes static quenching of the fluorophore (3,50), leading to dramatically reduced quantum yield, and to protein aggregation (Fig. 2A), highlighting the importance of preventing PP-IX aggregation to prevent porphyrin-mediated cell damage. Pharmacologic agents that affect PP-IX speciation could be therapeutically valuable for treating porphyria. Drugs targeting heme speciation are already known. For example, the antimalarial quinolone compounds prevent hemozoin granule formation by binding to heme monomers, and the heme monomer-quinolone complex is incorporated at the elongating hemozoin polymer, thereby inhibiting polymer elongation (51,52). Similarly, caffeine has been reported to prevent heme aggregation by forming a heme-caffeine monomeric complex (53). On the other hand, PP-IX and other related porphyrins are currently used as anti-tumor agents in photo-dynamic therapy (PDT). In the context of PDT, porphyrin-mediated protein aggregation and proteotoxicity play a beneficial cytotoxic role. For example, verteporfin a PDT agent, causes tumor selective protein aggregation (38). Thus, modulation of PP-IX speciation in a context-dependent manner could represent an attractive strategy to attenuate or enhance its toxicity.

### Selectivity and specificity of PP-IX-protein interaction and related protein aggregation is guided by PP-IX speciation

A hallmark of PP-IX induced protein aggregation is its selectivity. Some known PP-IX binding proteins, such as translocator protein (TSPO) and fatty acid binding protein 1 (FABP1), do not aggregate (39). Herein we observed that BSA does not bind PP-IX monomers (Fig. 4). Thus, we posit that PP-IX speciation might be one reason underlying the selectivity of PP-IX-protein interactions. PP-IX speciation also plays a role in the nature of protein aggregation. As we demonstrate (Fig. 3), detergent micellarized-PP-IX monomers lead to qualitatively and quantitatively different protein aggregates as compared to those formed by PP-IX dimers and higher order structures. Additionally, PP-IX mediated protein aggregation is organelle selective. Proteomic analysis of protein aggregation of ALA/DFO injected zebrafish livers showed aggregation of cytosolic, nuclear, ER and mitochondrial proteins but a total absence of lysosomal protein aggregation (37). Lysosomes accumulate porphyrins similar to the cytosol, mitochondria, ER and nucleus (54–57). However, lysosomes are acidic compartments with a luminal pH of 4.5-5 (58), which aligns well with our finding that PP-IX exists as higher order and relatively inert structures with lower oxidizing and protein aggregating ability at this pH. Furthermore,, porphyrin oligomers have lower quantum yield and impaired 1O_2_ generation capacity (59). Indeed, live cell imaging of ^1^O_2_ demonstrated that despite lysosomal and mitochondrial accumulation of porphyrin, ^1^O_2_ was generated only in the mitochondria (60). Thus, accumulation of anionic porphyrins such as PP-IX, in the acidic environment of lysosomes (57), is not accompanied by protein aggregation. In addition to higher order structure speciation, PP-IX monomerization also modulates its toxicity as shown for detergent monomerized PP-IX which has a higher oxidizing potential (Figs. 2,3). In liver, bile salts act as surfactant detergents, and by virtue of their ability to solubilize PP-IX, play an important role in porphyrin clearance. In fact, several mouse models of protoporphyria show the important role of bile salt-PP-IX interaction in the development of cholangiopathy (61,62).

### Impact of the albumin-PP-IX interaction

Albumin constitutes ~50 % of total plasma protein content (63). It is synthesized in hepatocytes then secreted into plasma (63,64) where it maintains oncotic pressure (65). In addition, albumin performs diverse functions ranging from ligand transport to serving as an antioxidant (63–66). Our data show that PP-IX binding to BSA leads to BSA oxidation and a conformational change (Figs. 5,7). Since albumin also binds to a vast network of proteins in the plasma, the so-called ‘albuminome’ (67–69), PP-IX binding could lead to disruption of components in the ‘albuminome’. In fact, oxidized human serum albumin is reported to differ significantly as compared with its un-oxidized counterpart with respect to ligand binding and antioxidant properties (70).

### Porphyrin-binding proteins as a PP-IX speciation agent

Secondary structure of albumin shows considerable Binding to albumin changes the toxicity of PP-IX’s by enhancing its oxidizing ability by shifting the equilibrium from higher order to dimeric PP-IX (Fig.8,10). On the other hand, albumin acts as a ‘PP-IX sponge’, and decreases endogenous PP-IX accumulation (11–13) and protein aggregation (Fig. 13). Our findings raise the question whether albumin modulates PP-IX induced protein aggregation intracellularly in hepatocytes or extracellularly in serum. Thus, by modulating speciation, PP-IX binding proteins such as albumin, might act as novel genetic modifiers of porphyric tissue damage.

## Conclusions

The reversible nature of PP-IX speciation and their different functional properties provide a potential target for therapeutic interventions. For example, our study demonstrates that PP-IX-induced protein aggregation could be reversed simply by a pH-induced shift in PP-IX speciation. This implies that transport of PP-IX-protein aggregates to the acidic lumen of the autophagolysosome can shift the equilibrium to higher order structures and lead to disaggregation of the PP-IX bound protein aggregates. Our findings highlight the dramatic difference in property and behavior of PP-IX, depending on its degree of oligomerization. Thus, PP-IX speciation could be utilized for designing better therapeutics for porphyria and improved PDT agents.

## EXPERIMENTAL PROCEDURES

### Cell lines and reagents

Human hepatocellular carcinoma cell line, HuH-7 cells (originally from the Japanese Collection of Research Bioresources Cell Bank) was a kind gift from Dr. Lei Yin (University of Michigan). HuH-7 cells were grown in Dulbecco’s modified Eagle’s medium (Cellgro, Manassas, VA), supplemented with 10% fetal bovine serum (Sigma-Aldrich).

PP-IX, deferoxamine mesylate (DFO), 5-ALA hydrochloride, N,N-dimethylacetamide (DMA), NaH_2_PO_4_, Na_2_HPO_4_, Flavin mono nucleotide (FMN), and ALA were obtained from Sigma Aldrich (St. Louis, MO). Fatty acid free bovine serum albumin (BSA) was from Calbiochem (San Diego, CA), coproporphyrin-III dihydrochloride (Copro) and uroporphyrin-III dihydrochloride (Uro) were obtained from Frontier Scientific (Logan UT).

### Absorbance/fluorescence measurements

PP-IX absorbance and fluorescence measurements were made using a TECAN SAFIRE II microplate reader running XFLUOR4SAFIREII (Version: V 4.62n), or the Spectramax id3 running SoftMax Pro.

### Quantum yield calculation

Absorbance (from 300-700 nm) and fluorescence emission spectra of 50 μM of PP-IX solution were collected. Relative quantum yield (Φ) were calculated using (Φ = ∫ *Em*/*Abs*),where ∫Em is the area under the curve for the emission spectra, and Abs is the absorbance at the excitation wavelength. ∫Em was calculated by integrating the emission spectrum using GraphPad Prismsoftware (GraphPad Software, San Diego, CA). The following excitation and emission wavelengths were used: Excitation - 437 nm, Emission - 460-850 nm (pH 4.5); Excitation - 368 nm, Emission - 400-850 nm (pH 7.4); Excitation - 389 nm, Emission - 410-850 nm (pH 9); Excitation - 410 nm, Emission - 450-850 nm (1% Empigen in pH 7.4).

### FMN Oxidation

An FMN stock solution (2.2 mM) was freshly prepared in 5 mM phosphate buffer (PB), pH 7.4 and stored away from light, on ice. The FMN stock solution was diluted to a final concentration of 22 μM in 400 mM of either NaH_2_PO_4_ (pH 4.5), PB (pH 7.4), Na_2_HPO_4_ (pH 9.0), or 1% Empigen BB in PB (pH 7.4), and incubated with different concentrations of PP-IX. DMA in all the reaction mixtures was maintained at 2.8% (v/v). The control reaction mixture contained FMN (22 μM) in 2.8% DMA in pH 4.5, 7.4, or 9 buffers. After incubating the reaction mixtures (30 min, ambient light), the fluorescence emission between 500-700 nm was recorded using a SAFIRE II plate reader, after exciting at 400 nm. The percent oxidation of FMN was estimated from the loss of FMN fluorescence at 533 nm after PP-IX treatment. Additionally, where indicated, PP-IX mediated FMN oxidation was assayed as above, but in the presence of the indicated concentrations of fatty acid-free BSA. The extent of FMN oxidation was estimated by the loss of absorbance at 370 and 450 nm.

### Treatment of HuH-7 cell lysates with PP-IX and biochemical analysis

Detergent-free HuH-7 cell lysates were prepared by 5 cycles of freeze-thawing then using a dounce homogenizer in 5 mM PB (pH 7.4) containing protease inhibitors cocktail (ThermoScientific, Waltham MA). The resulting solution was centrifuged (14,000 × g, 10 min, 4°C) and the supernatant and pellet were separated. The pellet was resuspended in phosphate buffered saline (PBS) supplemented with 2% SDS. Protein content in the supernatant and pellet fractions was determined using a bicinchoninic acid (BCA) assay. The final reaction mixture contained: 1 mg/ml protein with or without 50 μM PP-IX in 400 mM of either NaH_2_PO_4_ (pH 4.5), PB (pH 7.4), Na_2_HPO_4_ (pH 9.0), or 1% Empigen BB in PB (pH 7.4). Control samples had the same concentration of the vehicle, DMA. After incubating (30 min, 37°C, ambient light), the reaction was stopped by the addition of Laemmli sample buffer. Proteins (10 μg/condition) were separated by SDS-PAGE and stained with Coomassie blue or transferred to polyvinylidene fluoride (PVDF) membranes for immunoblotting. Immunoblots were visualized using horseradish peroxidase–tagged secondary antibody and chemiluminescence (Clarity Western ECL Substrate; BioRad, Hercules, CA). The antibodies used included those directed to lamin A/C, ubiquitin (Santa Cruz Biotechnology, Inc; Dallas, TX); lamin B1, p62 (Abcam; Cambridge, MA); K8 (clone TS1) and K18 (clone DC10), protein disulfide isomerase and BiP (Cell Signaling Technology; Danvers, MA).

### Calculation of number of PP-IX binding site on BSA and BSA-PP-IX dissociation constant

Fatty acid-free BSA (0.5 μM) was incubated with increasing concentrations of PP-IX (0-50 μM), at pH 4.5, 7.4, and 9 buffers, respectively, for 30 minutes. Then, the solution was excited (280 nm) and the fluorescence emission spectra were collected from 320-420 nm.

Data were analyzed using a double log plot of 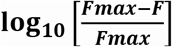 versus log_10_[*PP - IX*], where F_max_ and F are the fluorescence of BSA (347 nm) in the absence and presence of PP-IX (71). The plot yielded a straight line, and the number of binding sites was calculated using:

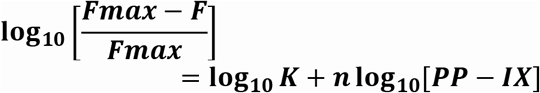

where, *K* is the binding constant, and *n* is the number of binding sites.

The dissociation constant (K_D_) and Hill coefficient (*h*) were calculated by plotting F_max_-F versus [PP-IX], and fitting the data to ‘Specific binding with Hill slope’ equation in GraphPad Prism. For n, *K*_D_, and h calculations, traces from each experiment were fitted individually and then averaged.

### Mass spectrometric analysis of PP-IX-mediated BSA oxidation

BSA (10 μM) was treated with 500 μM PP-IX in pH 4.5, 7.4 and 9 buffers (30 min, 22°C). After treatment, 100 μl of the reaction mixture was extracted by mixing with 850 μl of methanol and 50 μl of 0.1 N perchloric acid. The resulting suspension was pelleted (20,800×g, 4°C, 20 min). The pellet was solubilized in 8M guanidine hydrochloride, 50 mM Tris HCl, pH 8.1, 20 mM DTT and incubated at 60°C, 30 min to reduce disulfide bond, followed by incubation in the dark at room temperature with 40 mM of iodoacetamide to block the reduced cysteine residues. A fraction of the sample (10%) was desalted using ziptip C4 (EMD Millipore, Burlington, MA) and dried under vacuum. The sample was treated with 0.2 μg trypsin in 10 μl of 50 mM NH_4_HCO_3_ and incubated overnight at 37°C. The digested peptides were acidified with 1 μL of 10% formic acid followed by analysis using liquid chromatography-tandem mass spectrometry (LC-MS/MS). Samples were analyzed with a Q Exactive HF tandem mass spectrometer coupled to a Dionex Ultimate 3000 RLSCnano System (Thermo Scientific) by loading onto a fused silica trap column Acclaim PepMap 100, 75 μm x 2 cm (ThermoFisher). After washing (5 min, 5 μL/min with 0.1% TFA), the trap column was brought in-line with an analytical column (Nanoease MZ peptide BEH C18, 130Å, 1.7μm, 75 μm x 250 mm, Waters) for LC-MS/MS analysis. Peptides were eluted using a segmented linear gradient from 4 to 90% B (A: 0.2% formic acid, B: 0.08% formic acid, 80% acetonitrile): 4–15% B in 5min, 15-50% B in 50 min, and 50-90% B in 15 min. Mass spectrometry data was acquired using data-dependent acquisition with a cyclic series of a full scan from 250-2000 with resolution of 120,000 followed by MS/MS (HCD, relative collision energy 27%) of the 20 most intense ions (charge +1 to +6) and a dynamic exclusion duration of 20 sec. For database search, the peak list of the LC-MS/MS was generated by Thermo Proteome Discoverer (v. 2.1) into MASCOT Generic Format (MGF) and searched against the Uniprot bovine database plus a database composed of common laboratory contaminants using an in-house version of X!Tandem (72). The search parameters were: fragment mass error (20 ppm), parent mass error (+/-7ppm), fixed modification:C-carbamidolethyl); flexible modifications: Met oxidation for the primary search and other modifications were done at the refinement stage and less than 3 modifications are set for each search. Protease specificity: trypsin (miss 2 cuts), and only spectra with log_e_ <−2 were included in the final report. A detailed list of the oxidative modifications is listed in Supplementary Table S1.

### Treatment of HuH-7 cells with ALA/DFO ± BSA

HuH-7 cells were seeded in 10 cm dishes in complete (serum containing) media. When the cells reached confluency, the media was replaced with serum-free media with ALA/DFO (1 mM/100 μM) ± fatty acid-free BSA (5 mg/ml). Control dishes had serum-free media ± fatty acid-free BSA (5 mg/ml). After 16 h, the entire volume of medium was collected and then pelleted (3000 ×g, 10 min) to remove loating cells and debris. The supernatant was saved for analysis. For the remaining adherent cells, the culture plates were rinsed twice with PBS, and the cells were harvested by scraping in 10 ml PBS. The resulting cell suspension was centrifuged (400 × g, 5 min) followed by lysis of the cell pellet with PBS supplemented with 2% SDS then analysis by SDS-PAGE. The cell lysate and culture medium were also analyzed for porphyrin content as described below.

### Ultra Performance Liquid Chromatography (UPLC) analysis of porphyrins

A Waters ACQUITY UPLC system equipped with Empower software, ACQUITY H-Class PLUS (CH-A) Core, comprising of Quaternary Solvent Manager (QSM), a Sample Manager with Flow-Through Needle (SM-FTN), ACQUITY UPLC PDA Detector, ACQUITY UPLC Fluorescence Detector. A reverse-phase octadecylsilica (C18) ACQUITY UPLC BEH Shield RP18 Column, 130Å, 1.7μm, 3mm X 100mm UPLC column and ACQUITY UPLC BEH Shield RP18 VanGuard Pre-column, 130Å, 1.7μm, 2.1mm X 5mm was used. The fluorescence detector was set for excitation at 400 nm and recording emission at 635 nm. The column was eluted at a flow rate of 0.5 ml/min with linear gradients of solvents A and B (A, 0.1% formic acid in water; B, 0.1% formic acid in methanol). The solvent gradient was as follows: 0 to 1 min, 50-50% B; 1 to 2 min, 50-85% B; 2 to 3 min, 85-90% B; 3 to 9 min, 90-95% B; 9 to 9.5 min, 95-50% B; 9.5 to 14.5min, 50-50%. The cells were lysed after ALA+DFO treatment in PBS supplemented with 2% SDS. To extract porphyrins, DMA and 0.1% formic acid in methanol were added to the cell lysate or the isolated culture medium in a 1:1:2 ratio. After mixing then centrifuging (20,800 × g, 5 min) to precipitate proteins, 50 μl of the supernatant was injected into the UPLC system. Retention times of PP-IX, Uro and Copro were determined using commercially obtained standards.

### Porphyrin solution preparation

PP-IX was dissolved in N,N-dimethylacetamide (1 mg/ml), while Uro and Copro were dissolved in 100 mM NaOH, to prepare 5 mM stock solutions. Care was taken not to expose the stock solution to light. The stock solutions were diluted in appropriate buffers immediately prior to use. ALA and DFO were dissolved in water to yield 500 mM and 50 mM stock solutions, respectively, then filter-sterilized, aliquoted and stored at −20°C.

### Statistical Analysis

Statistical analysis of the quantum yield for different PP-IX nanostructures, FMN oxidation and porphyrin metabolite accumulation were performed using GraphPad Prism software (GraphPad Software; San Diego, CA). Statistical comparisons were carried out using ordinary one-way ANOVA analysis and Tukey’s multiple comparison test, or the unpaired T-test (2-tailed).

## Author Contributions

Conceptualization (DM), Formal analysis (DM, MBO), Funding acquisition (MBO, RB, BMP), Investigation (DM, BMP, AS, HZ), Methodology (DM, HZ), Project Administration (MBO), Resources (RB), Supervision (MBO), Visualization (DM), Writing – original draft (DM), Writing – review & editing (DM, RB, MBO).

## ACKNOWLEDGEMENTS

This work was supported by National Institutes of Health (NIH) R01 grants DK116548 (MBO) and GM130183 (RB); Honors Summer Fellowship (College of Literature, Science, and the Arts), and Summer Undergraduate Research Fellowship (Department of Molecular and Integrative Physiology) at the University of Michigan (BMP).

## Author Disclosure Statement

No competing financial interest exists.

## ABBREVIATION LIST

The abbreviations used are:

BSA: bovine serum albumin
Copro: coproporphyrin
DMA: N,N-dimethylacetamide
1O_2_: singlet oxygen
PP-IX: Protoporphyrin-IX
Φ: quantum yield
UPLC: Ultra Performance Liquid Chromatography
Uro: Uroporphyrin
UV-Vis: Ultra Violet-Visual spectrophotometry

## Supplementary Figures and Tables

**Supplementary Figure S1:**
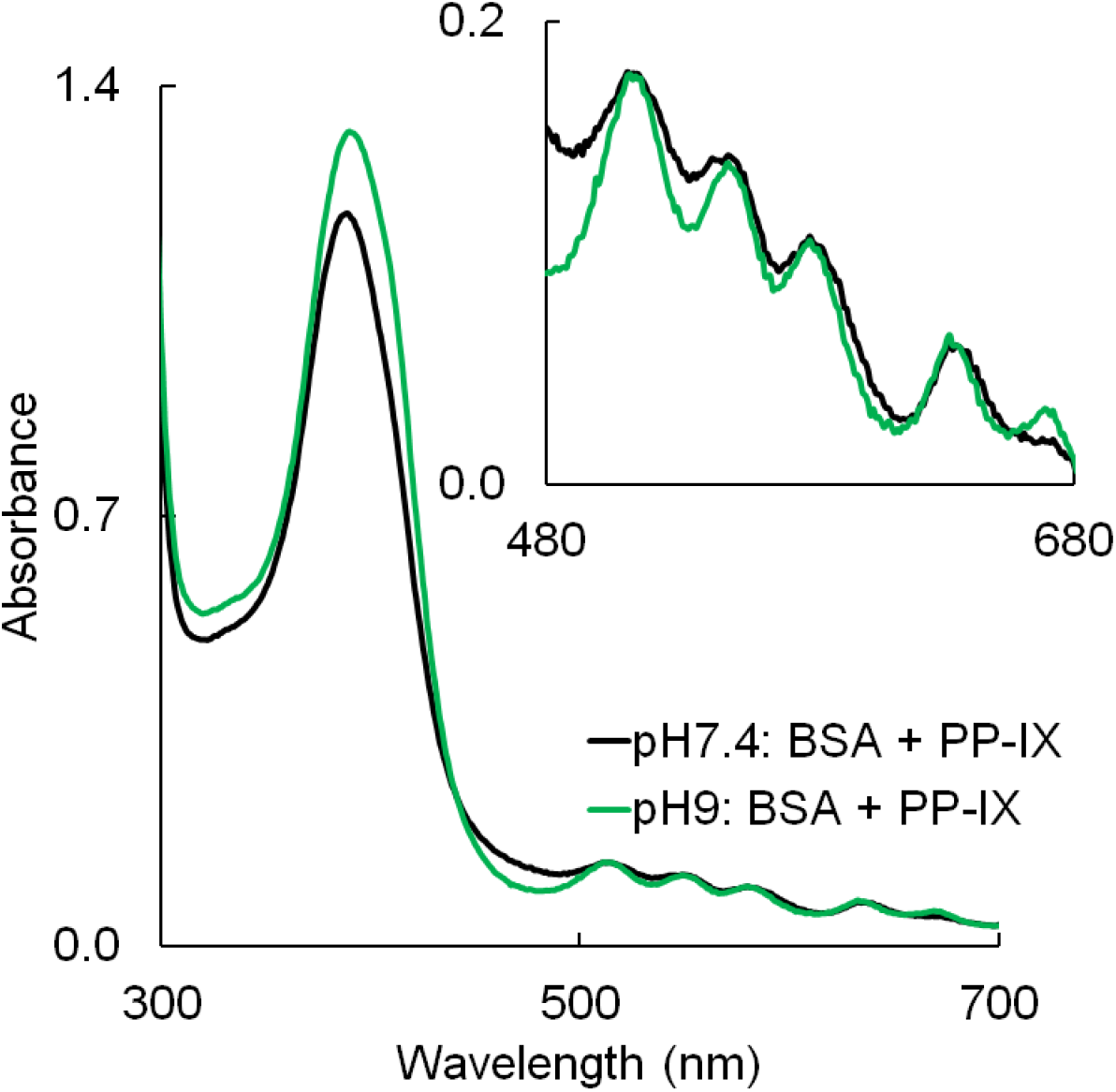
Overlay of BSA+PP-IX absorbance spectra collected at pH7.4 and pH9 (from the experiment described in Fig.4). Inset shows the zoomed part of the spectra from 480-680 nm. Data is representative of three independent experiments.

**Supplementary Table S1:** List of potential oxidative modifications on BSA that was searched for in PP-IX treated BSA sample.

| **Unimod**<br>**Accession**<br>**#** | **Interim Name** | **Description** | **Modified**<br>**amino acid** | **$\Delta$**<br>**monoisotopic**<br>**mass** |
| --- | --- | --- | --- | --- |
| 344 | Argglutamicsealde | Arginine oxidation to glutamic semialdehyde | R | -43 |
| 1914 | Met->AspSA | Methionine oxidation to aspartic semialdehyde | M | -32 |
| 360 | Pyrrolidinone | Proline oxidation to pyrrolidinone | P | -30 |
| 348 | His2Asn | His->Asn substitution | H | -23 |
| 349 | His2Asp | His->Asp substitution | H | -22 |
| 352 | Lysaminoadipicsealde | Lysine oxidation to aminoadipic semialdehyde | K | -1 |
| 351 | kynurenin | tryptophan oxidation to kynurenin | W | +4 |
| 359 | Pyroglutamic | proline oxidation to pyroglutamic acid | P | +14 |
| 288 | oxolactone | Tryptophan oxidation to oxolactone | W | +14 |
| 35 | Hydroxylation | Oxidation or Hydroxylation | MWY | +16 |
| 1922 | Pro->HAVA | Proline oxidation to 5-hydroxy-2-aminovaleric acid | P | +18 |
| 350 | hydroxykynurenin | tryptophan oxidation to hydroxykynurenin | W | +20 |
| 1923 | Delta:H(-4)O(2) | Tryptophan oxidation to beta-unsaturated-2,4-bis-tryptophandione | W | +28 |
| 425 | dihydroxy | dihydroxy | MWY | +32 |
| 1384 | Homocysteic_acid | methionine oxidation to homocysteic acid | M | +34 |
| 345 | Cysteic_acid | cysteine oxidation to cysteic acid | CWY | +48 |
| 1918 | Carbonyl | aldehyde and ketone modifications | AEILQRSV | +14 |

**Supplementary Table S2:** Table showing p-values from ordinary one-way ANOVA analysis and Tukey’s multiple comparison test for data shown in Fig. 2A.

| Tukey's multiple comparisons test | Significant? | Summary | Adjusted P Value |
| --- | --- | --- | --- |
| pH4.5 vs. pH7.4 | No | ns | 0.9997 |
| pH4.5 vs. pH9 | Yes | * | 0.0260 |
| pH4.5 vs. pH7.4 + Emp | Yes | **** | <0.0001 |
| pH7.4 vs. pH9 | Yes | * | 0.0294 |
| pH7.4 vs. pH7.4 + Emp | Yes | **** | <0.0001 |
| pH9 vs. pH7.4 + Emp | Yes | **** | <0.0001 |

**Table. S3:** The table shows p-values from ordinary one-way ANOVA analysis and Tukey’s multiple comparison test for data shown in Fig. 2C.

[PP-IX], 6.25 μM
| Tukey's multiple comparisons test | Significant? | Summary | Adjusted P Value |
| --- | --- | --- | --- |
| pH4.5 vs. pH7.4 | No | ns | >0.9999 |
| pH4.5 vs. pH9 | Yes | * | 0.0462 |
| pH4.5 vs. pH7.4 + Emp | No | ns | 0.0525 |
| pH7.4 vs. pH9 | Yes | * | 0.0463 |
| pH7.4 vs. pH7.4 + Emp | No | ns | 0.0526 |
| pH9 vs. pH7.4 + Emp | No | ns | 0.9997 |

[PP-IX], 12.5 μM
| Tukey's multiple comparisons test | Significant? | Summary | Adjusted P Value |
| --- | --- | --- | --- |
| pH4.5 vs. pH7.4 | No | ns | 0.3892 |
| pH4.5 vs. pH9 | Yes | *** | 0.0008 |
| pH4.5 vs. pH7.4 + Emp | Yes | *** | 0.0004 |
| pH7.4 vs. pH9 | Yes | ** | 0.0057 |
| pH7.4 vs. pH7.4 + Emp | Yes | ** | 0.0026 |
| pH9 vs. pH7.4 + Emp | No | ns | 0.9047 |

[PP-IX], 25 μM
| Tukey's multiple comparisons test | Significant? | Summary | Adjusted P Value |
| --- | --- | --- | --- |
| pH4.5 vs. pH7.4 | No | ns | 0.8757 |
| pH4.5 vs. pH9 | Yes | **** | <0.0001 |
| pH4.5 vs. pH7.4 + Emp | Yes | **** | <0.0001 |
| pH7.4 vs. pH9 | Yes | **** | <0.0001 |
| pH7.4 vs. pH7.4 + Emp | Yes | **** | <0.0001 |
| pH9 vs. pH7.4 + Emp | No | ns | 0.0998 |

[PP-IX], 30 μM
| Tukey's multiple comparisons test | Significant? | Summary | Adjusted P Value |
| --- | --- | --- | --- |
| pH4.5 vs. pH7.4 | No | ns | 0.9979 |
| pH4.5 vs. pH9 | Yes | *** | 0.0002 |
| pH4.5 vs. pH7.4 + Emp | Yes | **** | <0.0001 |
| pH7.4 vs. pH9 | Yes | *** | 0.0001 |
| pH7.4 vs. pH7.4 + Emp | Yes | **** | <0.0001 |
| pH9 vs. pH7.4 + Emp | Yes | * | 0.0131 |

[PP-IX], 30 μM
| Tukey's multiple comparisons test | Significant? | Summary | Adjusted P Value |
| --- | --- | --- | --- |
| pH4.5 vs. pH7.4 | No | ns | 0.2773 |
| pH4.5 vs. pH9 | Yes | **** | <0.0001 |
| pH4.5 vs. pH7.4 + Emp | Yes | **** | <0.0001 |
| pH7.4 vs. pH9 | Yes | **** | <0.0001 |
| pH7.4 vs. pH7.4 + Emp | Yes | **** | <0.0001 |
| pH9 vs. pH7.4 + Emp | Yes | *** | 0.0003 |

## REFERENCES

1. Dayan, F. E., and Dayan, E. A. (2011) Porphyrins: One Ring in the Colors of Life: A class of pigment molecules binds King George III, vampires and herbicides. American Scientist 99, 236–243

2. Gotardo, F., Cocca, L. H. Z., Acunha, T. V., Longoni, A., Toldo, J., Gonçalves, P. F. B., Iglesias, B. A., and De Boni, L. (2017) Investigating the intersystem crossing rate and triplet quantum yield of Protoporphyrin IX by means of pulse train fluorescence technique. Chemical Physics Letters 674, 48–57

3. Scolaro, L. M., Castriciano, M., Romeo, A., Patanè, S., Cefalì, E., and Allegrini, M. (2002) Aggregation Behavior of Protoporphyrin IX in Aqueous Solutions: Clear Evidence of Vesicle Formation. The Journal of Physical Chemistry B 106, 2453–2459

4. Isamu, I., and Kazutomo, U. (1991) Association Behavior of Protoporphyrin IX in Water and Aqueous Poly(N-vinylpyrrolidone) Solutions. Interaction between Protoporphyrin IX and Poly(N-vinylpyrrolidone). Bulletin of the Chemical Society of Japan 64, 2005–2007

5. Margalit, R., Shaklai, N., and Cohen, S. (1983) Fluorimetric studies on the dimerization equilibrium of protoporphyrin IX and its haemato derivative. Biochem J 209, 547–552

6. Seo, J., Jang, J., Warnke, S., Gewinner, S., Schöllkopf, W., and von Helden, G. (2016) Stacking Geometries of Early Protoporphyrin IX Aggregates Revealed by Gas-Phase Infrared Spectroscopy. Journal of the American Chemical Society 138, 16315–16321

7. Hunter, C. A., and Sanders, J. K. M. (1990) The nature of.pi.-.pi. interactions. Journal of the American Chemical Society 112, 5525–5534

8. Zhou, J., Li, J., Du, X., and Xu, B. (2017) Supramolecular biofunctional materials. Biomaterials 129, 1–27

9. Lehn, J. (1993) Supramolecular chemistry. Science 260, 1762–1763

10. Magna, G., Monti, D., Di Natale, C., Paolesse, R., and Stefanelli, M. (2019) The Assembly of Porphyrin Systems in Well-Defined Nanostructures: An Update. Molecules 24

11. Elemans, J. A. A. W., van Hameren, R., Nolte, R. J. M., and Rowan, A. E. (2006) Molecular Materials by Self-Assembly of Porphyrins, Phthalocyanines, and Perylenes. Advanced Materials 18, 1251–1266

12. Wurthner, F., Kaiser, T. E., and Saha-Moller, C. R. (2011) J-aggregates: from serendipitous discovery to supramolecular engineering of functional dye materials. Angew Chem Int Ed Engl 50, 3376–3410

13. Jin, C. S., Lovell, J. F., Chen, J., and Zheng, G. (2013) Ablation of Hypoxic Tumors with Dose-Equivalent Photothermal, but Not Photodynamic, Therapy Using a Nanostructured Porphyrin Assembly. ACS Nano 7, 2541–2550

14. Constantin, C., Neagu, M., Ion, R. M., Gherghiceanu, M., and Stavaru, C. (2010) Fullerene-porphyrin nanostructures in photodynamic therapy. Nanomedicine (Lond) 5, 307–317

15. Jin, C. S., Cui, L., Wang, F., Chen, J., and Zheng, G. (2014) Targeting-Triggered Porphysome Nanostructure Disruption for Activatable Photodynamic Therapy. Advanced Healthcare Materials 3, 1240–1249

16. Imahori, H., and Fukuzumi, S. (2004) Porphyrin- and Fullerene-Based Molecular Photovoltaic Devices. Advanced Functional Materials 14, 525–536

17. Hasobe, T., Imahori, H., Kamat, P. V., Ahn, T. K., Kim, S. K., Kim, D., Fujimoto, A., Hirakawa, T., and Fukuzumi, S. (2005) Photovoltaic Cells Using Composite Nanoclusters of Porphyrins and Fullerenes with Gold Nanoparticles. Journal of the American Chemical Society 127, 1216–1228

18. Walter, M. G., Rudine, A. B., and Wamser, C. C. (2010) Porphyrins and phthalocyanines in solar photovoltaic cells. Journal of Porphyrins and Phthalocyanines 14, 759–792

19. Zhang, N., Wang, L., Wang, H., Cao, R., Wang, J., Bai, F., and Fan, H. (2018) Self-Assembled One-Dimensional Porphyrin Nanostructures with Enhanced Photocatalytic Hydrogen Generation. Nano Letters 18, 560–566

20. Wang, Z., Ho, K. J., Medforth, C. J., and Shelnutt, J. A. (2006) Porphyrin Nanofiber Bundles from Phase-Transfer Ionic Self-Assembly and Their Photocatalytic Self-Metallization. Advanced Materials 18, 2557–2560

21. La, D. D., Bhosale, S. V., Jones, L. A., and Bhosale, S. V. (2017) Arginine-induced porphyrin-based self-assembled nanostructures for photocatalytic applications under simulated sunlight irradiation. Photochemical & Photobiological Sciences 16, 151–154

22. Chen, Y., Huang, Z.-H., Yue, M., and Kang, F. (2014) Integrating porphyrin nanoparticles into a 2D graphene matrix for free-standing nanohybrid films with enhanced visible-light photocatalytic activity. Nanoscale 6, 978–985

23. Medforth, C. J., Wang, Z., Martin, K. E., Song, Y., Jacobsen, J. L., and Shelnutt, J. A. (2009) Self-assembled porphyrin nanostructures. Chemical Communications, 7261–7277

24. Dini, F., Martinelli, E., Pomarico, G., Paolesse, R., Monti, D., Filippini, D., D’Amico, A., Lundstrom, I., and Di Natale, C. (2009) Chemical sensitivity of self-assembled porphyrin nano-aggregates. Nanotechnology 20, 055502

25. Wang, Z., Medforth, C. J., and Shelnutt, J. A. (2004) Porphyrin Nanotubes by Ionic Self-Assembly. Journal of the American Chemical Society 126, 15954–15955

26. Puy, H., Gouya, L., and Deybach, J.-C. (2010) Porphyrias. The Lancet 375, 924–937

27. Maitra, D., Bragazzi Cunha, J., Elenbaas, J. S., Bonkovsky, H. L., Shavit, J. A., and Omary, M. B. (2019) Porphyrin-Induced Protein Oxidation and Aggregation as a Mechanism of Porphyria-Associated Cell Injury. Cell Mol Gastroenterol Hepatol 8, 535–548

28. Schultz, I. J., Chen, C., Paw, B. H., and Hamza, I. (2010) Iron and porphyrin trafficking in heme biogenesis. J Biol Chem 285, 26753–26759

29. Montgomery Bissell, D. (2015) Chapter 66 - The Porphyrias. in Rosenberg’s Molecular and Genetic Basis of Neurological and Psychiatric Disease (Fifth Edition) (Rosenberg, R. N., and Pascual, J. M. eds.), Academic Press, Boston. pp 731–749

30. Ajioka, R. S., Phillips, J. D., and Kushner, J. P. (2006) Biosynthesis of heme in mammals. Biochim Biophys Acta 1763, 723–736

31. Badminton, M. N., and Elder, G. H. (2014) CHAPTER 28 - The porphyrias: inherited disorders of haem synthesis. in Clinical Biochemistry: Metabolic and Clinical Aspects (Third Edition) (Marshall, W. J., Lapsley, M., Day, A. P., and Ayling, R. M. eds.), Churchill Livingstone. pp 533–549

32. Foote, C. S. (1991) DEFINITION OF TYPE I and TYPE II PHOTOSENSITIZED OXIDATION. Photochemistry and Photobiology 54, 659–659

33. Takeshita, K., Takajo, T., Hirata, H., Ono, M., and Utsumi, H. (2004) In Vivo Oxygen Radical Generation in the Skin of the Protoporphyria Model Mouse with Visible Light Exposure: An L-Band ESR Study. Journal of Investigative Dermatology 122, 1463–1470

34. Brun, A., and Sandberg, S. (1991) Mechanisms of photosensitivity in porphyric patients with special emphasis on erythropoietic protoporphyria. Journal of Photochemistry and Photobiology B: Biology 10, 285–302

35. Singla, A., Griggs, N. W., Kwan, R., Snider, N. T., Maitra, D., Ernst, S. A., Herrmann, H., and Omary, M. B. (2013) Lamin aggregation is an early sensor of porphyria-induced liver injury. J Cell Sci 126, 3105–3112

36. Saggi, H., Maitra, D., Jiang, A., Zhang, R., Wang, P., Cornuet, P., Singh, S., Locker, J., Ma, X., Dailey, H., Abrams, M., Omary, M. B., Monga, S. P., and Nejak-Bowen, K. (2019) Loss of hepatocyte beta-catenin protects mice from experimental porphyria-associated liver injury. J Hepatol 70, 108–117

37. Elenbaas, J. S., Maitra, D., Liu, Y., Lentz, S. I., Nelson, B., Hoenerhoff, M. J., Shavit, J. A., and Omary, M. B. (2016) A precursor-inducible zebrafish model of acute protoporphyria with hepatic protein aggregation and multiorganelle stress. FASEB J 30, 1798–1810

38. Zhang, H., Ramakrishnan, S. K., Triner, D., Centofanti, B., Maitra, D., Gyorffy, B., Sebolt-Leopold, J. S., Dame, M. K., Varani, J., Brenner, D. E., Fearon, E. R., Omary, M. B., and Shah, Y. M. (2015) Tumor-selective proteotoxicity of verteporfin inhibits colon cancer progression independently of YAP1. Sci Signal 8, ra98

39. Maitra, D., Elenbaas, J. S., Whitesall, S. E., Basrur, V., D’Alecy, L. G., and Omary, M. B. (2015) Ambient Light Promotes Selective Subcellular Proteotoxicity after Endogenous and Exogenous Porphyrinogenic Stress. J Biol Chem 290, 23711–23724

40. Fernandez, N. F., Sansone, S., Mazzini, A., and Brancaleon, L. (2008) Irradiation of the Porphyrin Causes Unfolding of the Protein in the Protoporphyrin IX/β-Lactoglobulin Noncovalent Complex. The Journal of Physical Chemistry B 112, 7592–7600

41. Belcher, J., Sansone, S., Fernandez, N. F., Haskins, W. E., and Brancaleon, L. (2009) Photoinduced Unfolding of β-Lactoglobulin Mediated by a Water-Soluble Porphyrin. The Journal of Physical Chemistry B 113, 6020–6030

42. Maitra, D., Carter, E. L., Richardson, R., Rittie, L., Basrur, V., Zhang, H., Nesvizhskii, A. I., Osawa, Y., Wolf, M. W., Ragsdale, S. W., Lehnert, N., Herrmann, H., and Omary, M. B. (2019) Oxygen and Conformation Dependent Protein Oxidation and Aggregation by Porphyrins in Hepatocytes and Light-Exposed Cells. Cell Mol Gastroenterol Hepatol 8, 659–682 e651

43. Teng, K. W., and Lee, S. H. (2019) Characterization of Protoporphyrin IX Species in Vitro Using Fluorescence Spectroscopy and Polar Plot Analysis. The Journal of Physical Chemistry B 123, 5832–5840

44. Sulkowski, L., Pawelczak, B., Chudzik, M., and Maciazek-Jurczyk, M. (2016) Characteristics of the Protoporphyrin IX Binding Sites on Human Serum Albumin Using Molecular Docking. Molecules 21

45. Brancaleon, L., and Moseley, H. (2002) Effects of photoproducts on the binding properties of protoporphyrin IX to proteins. Biophysical Chemistry 96, 77–87

46. Shaw, A. K., and Pal, S. K. (2008) Resonance energy transfer and ligand binding studies on pH-induced folded states of human serum albumin. J Photochem Photobiol B 90, 187–197

47. Camilloni, C., Rocco, A. G., Eberini, I., Gianazza, E., Broglia, R. A., and Tiana, G. (2008) Urea and guanidinium chloride denature protein L in different ways in molecular dynamics simulations. Biophys J 94, 4654–4661

48. Baral, A., Satish, L., Das, D. P., Sahoo, H., and Ghosh, M. K. (2017) Construing the interactions between MnO2 nanoparticle and bovine serum albumin: insight into the structure and stability of a protein–nanoparticle complex. New Journal of Chemistry 41, 8130–8139

49. Wang, M., Wey, S., Zhang, Y., Ye, R., and Lee, A. S. (2009) Role of the unfolded protein response regulator GRP78/BiP in development, cancer, and neurological disorders. Antioxid Redox Signal 11, 2307–2316

50. Sansaloni-Pastor, S., Bouilloux, J., and Lange, N. (2019) The Dark Side: Photosensitizer Prodrugs. Pharmaceuticals (Basel) 12

51. Sullivan, D. J., Jr., Matile, H., Ridley, R. G., and Goldberg, D. E. (1998) A common mechanism for blockade of heme polymerization by antimalarial quinolines. J Biol Chem 273, 31103–31107

52. Sullivan, D. J., Jr., Gluzman, I. Y., Russell, D. G., and Goldberg, D. E. (1996) On the molecular mechanism of chloroquine’s antimalarial action. Proc Natl Acad Sci U S A 93, 11865–11870

53. Carter, E. L., Ramirez, Y., and Ragsdale, S. W. (2017) The heme-regulatory motif of nuclear receptor Rev-erbbeta is a key mediator of heme and redox signaling in circadian rhythm maintenance and metabolism. J Biol Chem 292, 11280–11299

54. Ezzeddine, R., Al-Banaw, A., Tovmasyan, A., Craik, J. D., Batinic-Haberle, I., and Benov, L. T. (2013) Effect of molecular characteristics on cellular uptake, subcellular localization, and phototoxicity of Zn(II) N-alkylpyridylporphyrins. J Biol Chem 288, 36579–36588

55. Hsieh, Y. J., Wu, C. C., Chang, C. J., and Yu, J. S. (2003) Subcellular localization of Photofrin determines the death phenotype of human epidermoid carcinoma A431 cells triggered by photodynamic therapy: when plasma membranes are the main targets. J Cell Physiol 194, 363–375

56. Teiten, M. H., Bezdetnaya, L., Morliere, P., Santus, R., and Guillemin, F. (2003) Endoplasmic reticulum and Golgi apparatus are the preferential sites of Foscan localisation in cultured tumour cells. Br J Cancer 88, 146–152

57. Woodburn, K. W., Vardaxis, N. J., Hill, J. S., Kaye, A. H., and Phillips, D. R. (1991) SUBCELLULAR LOCALIZATION OF PORPHYRINS USING CONFOCAL LASER SCANNING MICROSCOPY. Photochemistry and Photobiology 54, 725–732

58. Mindell, J. A. (2012) Lysosomal acidification mechanisms. Annu Rev Physiol 74, 69–86

59. Tanielian, C., Schweitzer, C., Mechin, R., and Wolff, C. (2001) Quantum yield of singlet oxygen production by monomeric and aggregated forms of hematoporphyrin derivative. Free Radical Biology and Medicine 30, 208–212

60. Kim, S., Tachikawa, T., Fujitsuka, M., and Majima, T. (2014) Far-Red Fluorescence Probe for Monitoring Singlet Oxygen during Photodynamic Therapy. Journal of the American Chemical Society 136, 11707–11715

61. Fickert, P., Stoger, U., Fuchsbichler, A., Moustafa, T., Marschall, H. U., Weiglein, A. H., Tsybrovskyy, O., Jaeschke, H., Zatloukal, K., Denk, H., and Trauner, M. (2007) A new xenobiotic-induced mouse model of sclerosing cholangitis and biliary fibrosis. Am J Pathol 171, 525–536

62. Lyoumi, S., Abitbol, M., Rainteau, D., Karim, Z., Bernex, F., Oustric, V., Millot, S., Letteron, P., Heming, N., Guillmot, L., Montagutelli, X., Berdeaux, G., Gouya, L., Poupon, R., Deybach, J. C., Beaumont, C., and Puy, H. (2011) Protoporphyrin retention in hepatocytes and Kupffer cells prevents sclerosing cholangitis in erythropoietic protoporphyria mouse model. Gastroenterology 141, 1509–1519, 1519 e1501-1503

63. Quinlan, G. J., Martin, G. S., and Evans, T. W. (2005) Albumin: Biochemical properties and therapeutic potential. Hepatology 41, 1211–1219

64. Rabbani, G., and Ahn, S. N. (2019) Structure, enzymatic activities, glycation and therapeutic potential of human serum albumin: A natural cargo. International Journal of Biological Macromolecules 123, 979–990

65. Evans, T. W. (2002) Review article: albumin as a drug—biological effects of albumin unrelated to oncotic pressure. Alimentary Pharmacology & Therapeutics 16, 6–11

66. Fasano, M., Curry, S., Terreno, E., Galliano, M., Fanali, G., Narciso, P., Notari, S., and Ascenzi, P. (2005) The extraordinary ligand binding properties of human serum albumin. IUBMB Life 57, 787–796

67. (2007) The Albuminome as a Tool for Biomarker Discovery. in Clinical Proteomics. pp 263–278

68. Liu, Z., Li, S., Wang, H., Tang, M., Zhou, M., Yu, J., Bai, S., Li, P., Zhou, J., and Xie, P. (2017) Proteomic and network analysis of human serum albuminome by integrated use of quick crosslinking and two-step precipitation. Sci Rep 7, 9856

69. Gundry, R. L., Fu, Q., Jelinek, C. A., Van Eyk, J. E., and Cotter, R. J. (2007) Investigation of an albumin-enriched fraction of human serum and its albuminome. Proteomics Clin Appl 1, 73–88

70. Kawakami, A., Kubota, K., Yamada, N., Tagami, U., Takehana, K., Sonaka, I., Suzuki, E., and Hirayama, K. (2006) Identification and characterization of oxidized human serum albumin. The FEBS Journal 273, 3346–3357

71. Sen, P., Fatima, S., Ahmad, B., and Khan, R. H. (2009) Interactions of thioflavin T with serum albumins: spectroscopic analyses. Spectrochim Acta A Mol Biomol Spectrosc 74, 94–99

72. Craig, R., and Beavis, R. C. (2004) TANDEM: matching proteins with tandem mass spectra. Bioinformatics 20, 1466–1467

